# RNA Structure Directs RNA Partitioning and is Actively Disrupted inside Stress Granules to Enable Cellular Recovery

**DOI:** 10.1101/2025.06.05.658090

**Authors:** Bowen Zhu, Jovi Jian An Lim, Yu Zhang, Wen Ting Tan, Jiayi Zhang, Joy S Xiang, Roland Huber, Anthony Khong, Yue Wan

## Abstract

RNA structures play important roles in liquid-liquid phase separation. However, how it is regulated during stress response and stress granule formation is still under studied. Here, we performed *in vivo* RNA structure probing before and after sodium arsenite treatment, and in stress granules. While RNAs generally become more double-stranded upon stress, they maintain their single-strandedness inside stress granules. We showed that RNA single-strandedness enables increased inclusion inside stress granules and that stress granule-enriched RNAs form fewer intra- and intermolecular RNA-RNA interactions. Additionally, several RNA binding proteins including SRSF1 are enriched in differential structure regions. eCLIP analysis revealed that SRSF1 binds to single-stranded regions along RNAs, and increased SRSF1 binding enabled better inclusion of RNAs in stress granules, whereas depletion of SRSF1 decreased stress granule formation under mild oxidative stress. We also observed the active unwinding of RNAs inside stress granules regulated by helicases, including DDX3X, and showed that inhibition of DDX3X results in slower dissolution of stress granules during recovery. Our study reveals the existence of multiple mechanisms to maintain RNA single-strandedness inside stress granules and to allow reversibility of stress granule formation, highlighting the importance of regulating RNA structure to enable cellular plasticity and stress response.

## Introduction

Efficient response to environmental stress is important for all organisms. Stress granules (SGs) are transient cytosolic messenger ribonucleoprotein (mRNP) biomolecular condensates that are induced by acute reduction in global protein synthesis while cells encounter adverse environments. In general, stress granules formation and dissolution are thought to play an adaptive role in helping cells survive under a variety of stressors including oxidative stress, hypoxia, ER-stress, and virus infection (Marcelo et al., 2021). Prolonged persistence of stress granules is thought to alter stress granule dynamics which promotes protein aggregation and neuronal degenerative diseases (Jain and Vale, 2017; Lavalee et al., 2021; Lee et al., 2013; Lloyd, 2013; Somasekharan et al., 2015; Song and Grabocka, 2023). Therefore, it is of significant interest to understand stress granule biology, how they are formed and dissolved, and how their dysregulation could contribute to diseases.

Mechanistically, stress granules form via the condensation of the accumulated non-translated mRNPs in response to ribosome release during cellular stress. The condensation is thought to be mediated by multivalent interactions via biomolecules such as G3BP1/2 and RNAs and is driven by liquid-liquid phase separation (Anderson et al., 2015; Lavalee et al., 2021; Protter and Parker, 2016; Wheeler et al., 2016). While significant progress has been achieved in understanding the key condensate proteins like G3BP1/2 in stress granule assembly (Guillen-Boixet et al., 2020; Jain et al., 2016; Sanders et al., 2020; Yang et al., 2020), the role of RNA in this process remains controversial (Van Treeck et al., 2018; Parker et al., 2024; Trussina et al., 2024; Glauninger et al., 2024). Recent studies have revealed RNA features such as AU content and transcript length have been shown to impact the ability of RNA to partition into stress granules (Khong et al., 2016; Namkoong et al., 2017), however, the role of RNA structures and RNA-RNA interactions in stress granule assembly is debated. As such, direct measurements of RNA structures and RNA-RNA pairing would be highly informative in helping to elucidate the role of RNA in stress granule condensation, RNA recruitment to stress granules, and stress granule function in modulating gene expression.

To address the aforementioned fundamental questions, we conducted the first comprehensive analysis of the “RNA structurome” under arsenite stress in whole cells and stress granules. Our findings reveal that while RNAs generally become more double-stranded during arsenite stress, stress granule-enriched RNAs remain largely single-stranded. Additionally, we observed that both intra- and intermolecular RNA-RNA interactions are decreased in stress granule-enriched RNAs and that they are actively unwound inside stress granules. We also identified different RNA binding proteins (RBPs) that are responsible for recruiting and unwinding RNAs inside stress granules and show that inhibition of one of these RBPs, DDX3X, inhibits stress granule dissolution during post-stress recovery. These results suggest that RNA structures play key roles in the reversible assembly and disassembly of cellular compartments to enable proper gene regulation in response to environmental changes.

## Results

### The cellular transcriptome becomes more double-stranded upon arsenite stress

To study dynamic changes in RNA structures and RNA-RNA interactions during arsenite stress, we performed *in vivo* click Selective 2-Hydroxyl Acylation and Profiling Experiment (icSHAPE) and proximity ligation sequencing (SPLASH, **Figure 1A**). Specifically, we stressed HeLa cells with sodium arsenite (NaAsO_2_) for an hour and then allowed the cells to recover for 1 or 2 hours by removing sodium arsenite from the media. At each of these time points (untreated, 1-hour arsenite stress, 1-hour recovery, and 2-hour recovery), we treated the cells with the SHAPE compound, NAI-N_3_, to map local secondary structures. As both intra- and inter-molecular RNA-RNA interactions are possibly involved in the process of biomolecular condensate formation, we performed SPLASH to determine the intra- and inter-molecular RNA-RNA interactions in untreated and stressed cells. Additionally, we also performed ribosome profiling (Ribo-seq) to study the translation efficiency of the cellular transcripts during stress and recovery (**Figure 1A**).

**Figure 1.**
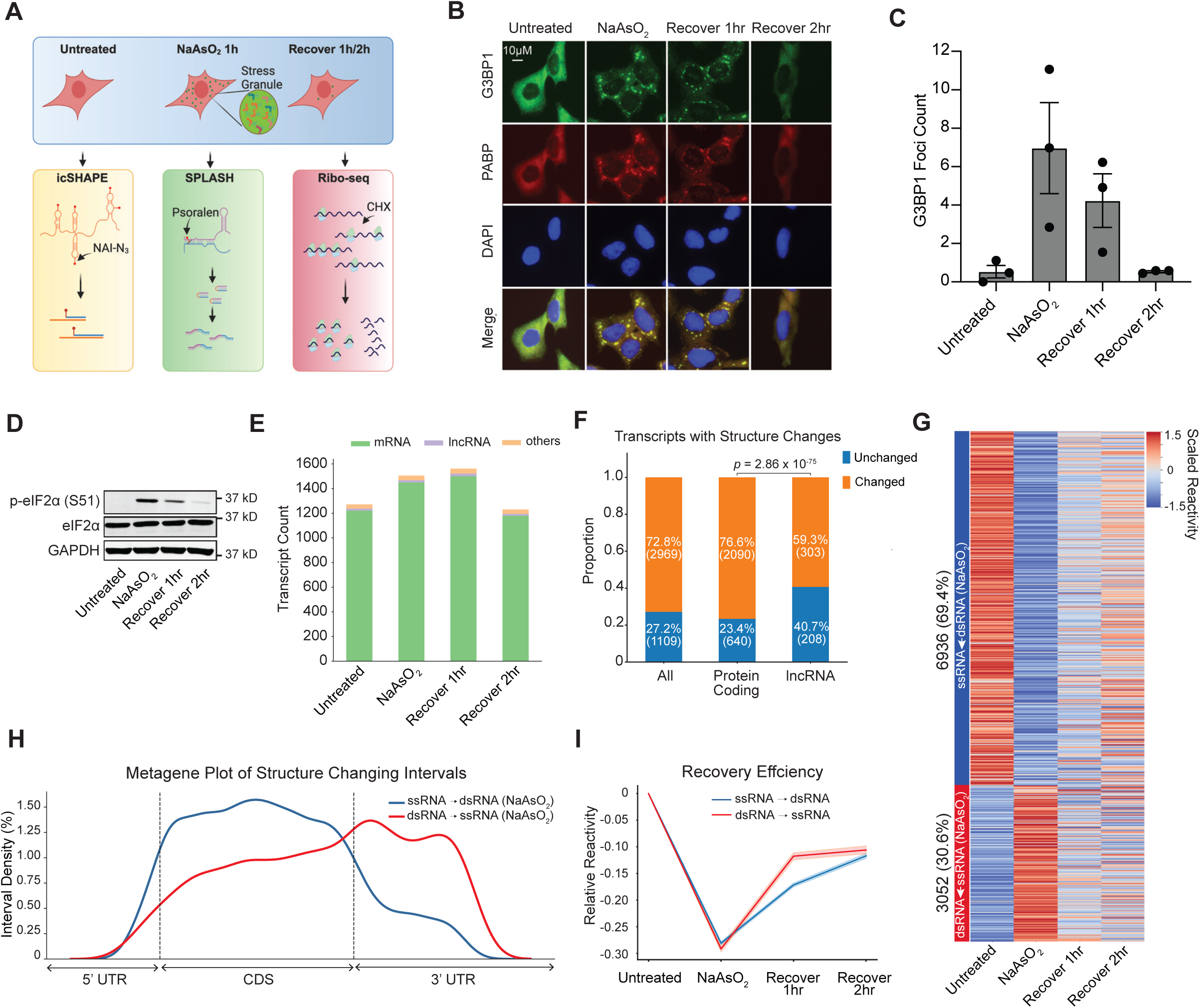
Sodium arsenite treatment induces global increase in RNA secondary structures. **A**, Schematic of the experimental workflow: icSHAPE, SPLASH, and Ribo-seq were performed to investigate the changes at the levels of RNA structure, RNA-RNA interaction, and mRNA translation. **B**, Immuno-fluorescence staining of G3BP1 (green), PABP (red), and nuclei (labeled as DAPI in blue) in Hela cells after treatment with sodium arsenite for 1 hour and recovered for 1 or 2 hours. **C**, Barplot showing the quantification of stress granule numbers formed in unstressed Hela cells, Hela cells that are treated with sodium arsenite, treated cells that are recovered for 1 hour, and for 2 hours, respectively. **D**, Western blot analysis of phospho-eIF2α (Ser-51), total eIF2α, and GAPDH in unstressed Hela cells, Hela cells that are treated with sodium arsenite, treated cells that are recovered for 1 hour, and for 2 hours, respectively. **E**, Stacked bar plot showing the distribution of biotypes of the genes detected in unstressed Hela cells, Hela cells that are treated with sodium arsenite, treated cells that are recovered for 1 hour, and for 2 hours, respectively. **F**, Stacked bar plot showing the proportion of changes in SHAPE reactivity upon arsenite-induced cellular stress all transcripts, protein coding genes, and lncRNAs. *p*-values were calculated using Chi-square Test. **G**, Heatmap showing reactivity in RNA regions in untreated cells, sodium arsenite treated cells, in treated cells that are recovered for 1 hour and in treated cells that are recovered for 2 hours. **H**, Density plots showing the change in reactivity of the nucleotides on indicated localizations of mRNA, namely 5’UTR, CDS and 3’UTR. **I**, Line plots showing the recovery rate of RNA regions that has become more double-stranded or more single-stranded upon sodium arsenite stress.

We first checked that treating the cells with sodium arsenite resulted in an increased number of G3BP1-positive stress granules, by performing immunofluorescence staining using the antibody against G3BP1, a well-characterized stress granule nucleator (**Figure 1B, 1C**). As expected, we discovered that the stress granules form after 1 hour of treatment with sodium arsenite and that the number of stress granules decreased as the cells recovered for 1 and 2 hours, with most of the stress granules disassembled by 2 hours post recovery (**Figure 1B, 1C**). Additionally, we also examined the phosphorylation of the translation initiation factor EIF2α upon stress and recovery. As expected, we observed a drastic induction of the phosphorylation of EIF2α at the Ser51 site 1-hour post sodium arsenite treatment, followed by a reduction at 1 and 2 hours after sodium arsenite removal (**Figure 1D**), confirming that our experimental setting indeed induced the integrated stress response, marked by reversible translational shutoff and stress granule formation.

To map the RNA structure of the stressed and recovered cellular transcriptome, we performed two biological replicates of icSHAPE on each of the four time points and evaluated its reproducibility, accuracy, and sequencing depth to ensure that we obtain high-quality data on RNA structure dynamic changes during stress. We discovered a high correlation of reverse transcriptase stop sites between biological replicates at each time point (*R*>0.8, **Supp. Figure 1A**), indicating that our structure probing is robust. In addition, mapping the reactivity of 28S rRNA to its known structure also showed a high AUC-ROC (AUC-ROC>0.7), indicating that our structure probing is accurate (**Supp. Figure 1B**). We detected RNA structure probing information for more than 1,200 genes at each time point (**Figure 1E**), among which 1,109 genes (1,069 protein-coding, 40 non-coding) are shared across the four time points, allowing us to compare structure between control, stressed, and recovery states for a significant fraction of the human transcriptome. In addition to RNA structure information, our icSHAPE experiments can also determine the transcript abundance of RNAs in each condition, as calculated from the reads in the DMSO libraries. We confirmed that most RNAs do not change their expression upon sodium arsenite stress, with ∼1.8% of detected transcripts showing gene expression changes (**Supp. Figure 1C**). As expected, GO term analysis showed that these transcripts are enriched in several stress-related pathways, including unfolded protein response, response to heat shock, and stress response to metal ions, indicating that our treatment and analysis are robust (**Supp. Figure 1D**).

Upon sodium arsenite treatment, we observed that ∼1% of the RNA structure probed bases changed structure and that these structure changes are widely distributed across most RNA transcripts (72.8%, **Supp. Figure 1E**). As compared to protein-coding mRNAs, long non-coding RNAs showed a lower proportion of structural changes (**Figure 1F**), suggesting that they are more structurally stable under stress. Interestingly, we observed that most of the RNAs become more double-stranded upon stress, with 69.4% of the structure-changing bases becoming more paired (**Figure 1G**). Metagene analysis of the structure changing intervals centered at the translational start and stop sites showed that the coding region generally becomes more paired, while the 3’UTR region becomes less paired upon stress (**Figure 1H**). This agrees with the hypothesis that translation is inhibited during sodium arsenite treatment and that local RNA structures are restored on the coding sequence when the ribosomes fall off (Rouskin et al., 2014). To gain insights into the RNA structure dynamics post-stress, we grouped structure-changing windows according to their stress-induced structure alteration and calculated the mean reactivity during stress and recovery. Interestingly, RNAs that had become more single-stranded upon stress recovered faster than the bases that had become more double-stranded during stress (**Figure 1I**). As double-stranded regions may require active unwinding processes to recover, we hypothesize that the need for helicases could be one reason that double-stranded regions recover slower than single-stranded regions.

To determine whether our observed structural changes are intrinsic to the RNA molecules or a result of the presence of other *in vivo* cellular factors, we performed a control experiment, whereby we extracted the RNAs from cells, refolded them, and then performed NAI-N_3_ labeling on the refolded transcriptome *in vitro*. As expected, we observed a high correlation (*R* >0.82) for RT-stops between two biological replicates at each time point (**Supp. Figure 2A**) and that *in vitro* refolded RNAs are more double-stranded than transcripts inside cells (**Supp. Figure 2B**). However, in contrast to our previous observation that RNAs become more paired during stress, we did not observe this increased pairing for *in vitro* refolded RNAs globally or locally (**Supp. Figure 2B, C**), indicating that the increased double-strandedness upon stress occurs mainly *in vivo*.

### RNAs remain single-stranded inside the stress granule upon arsenite stress

Knowing that we can reliably measure RNA-structure changes during arsenite stress and recovery and that it aligns with other published observations (Rouskin et al., 2014), we sought to ask whether the RNAs found within stress granules are structurally different from the RNAs outside of the stress granules. Specifically, we treated the cells with NAI-N_3_, enriched RNA granules via subcellular fractionation, and prepared sequencing libraries (hereafter referred to as Frac-SHAPE) from the enriched RNA (**Figure 2A, Methods**). We first confirmed that our stress granule enrichment is robust by detecting known stress granule-enriched and -depleted RNAs via quantitative RT-PCR (**Supp. Figure 3A**), and by showing that we obtained a good correlation with a published stress granule enrichment dataset (**Supp. Figure 3B**) (Khong et al., 2017). Interestingly, we discovered that RNAs inside the stress granule compartment are less paired than RNAs in the soluble cytoplasm (**Figure 2B**), which is surprising as the concentration of RNA inside stress granules is higher than that of RNAs inside the cytosol (Jain et al., 2016).

**Figure 2.**
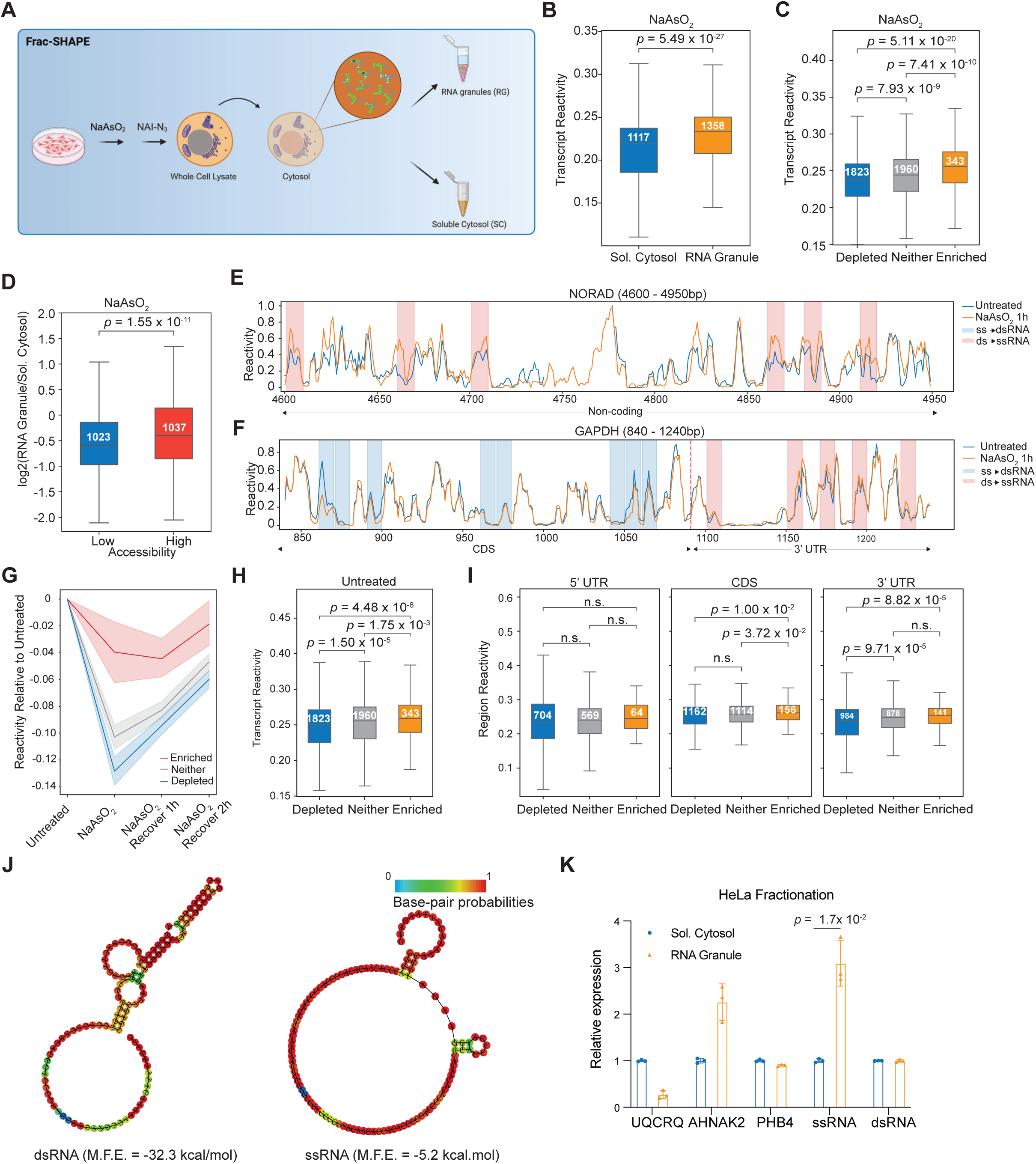
Stress granule-enriched RNAs are more single-stranded than cytosolic RNAs upon stress. **A**, Workflow of RNA structure probing in stress granules. We performed sodium arsenite treatment followed by subcellular fractionation to separate RNA granules (RG) from the soluble cytosol (SC). We then performed NAI-N_3_ treatment on the two fractions. **B**, Boxplot showing the distribution of reactivities of RNAs that are detected in the soluble cytosol and RNA granules. **C**, Boxplot showing the reactivities for stress-granule-enriched RNA, stress-granule-depleted-RNA, or neither transcripts after sodium arsenite treatment. **D**, Boxplot showing the distribution of RNA granule enrichment for the RNAs with high or low reactivity. Statistical analysis of **B-D** was performed using Mann-Whitney U Test, with sample sizes and p-values labelled in the panels. **E,F,** Line plots showing the transcript reactivity of a representative region of NORAD (stress-granule-enriched, **E**) and GAPDH RNA (stress-granule-depleted, **F**). Intervals in blue and red represent significant decrease and increase in single-strandedness respectively. **G**, Line plots showing the relative changes in reactivity for stress-granule-enriched, stress-granule-depleted, and neither transcripts, during sodium arsenite treatment and recovery. **H, I**, Boxplots showing the distribution of reactivity for stress-granule-enriched, stress-granule-depleted, and neither transcripts (**H**) and their indicated regions (**I**) in unstressed HeLa cells. Statistical analysis of **H,I** was done using Mann-Whitney U Test, with sample sizes and p-values labelled in the panels. **J**, Secondary structure models of two synthetic RNA molecules with identical length and GC content but with different propensities to form secondary structures, as predicted using Vienna RNA WebServer. We indicate the minimum free energy (M.F.E) and base-pair probabilities for each RNA. **K**, Bar plots showing the extent of RNA granule enrichment for the single-stranded and the double-stranded RNA. UQCRQ (depleted), AHNAK2 (enriched), and PHB4 (neither) were used as controls. Statistical analysis of **K** was performed using the Student’s T-Test. Data are shown as mean ± SEM, n = 3.

To confirm this notion, we separated transcripts according to their localization, namely whether they are enriched, depleted, or present at similar levels (neither) in stress granules, as compared to inside the cytoplasm (**Methods**). As previously described in the literature, stress granule-enriched RNAs are lower in GC content (more AU-rich) (**Supp. Figure 3C**) and are longer (**Supp. Figure 3D**). We further confirmed that our stress granule-enriched RNAs were translationally repressed by performing ribosome profiling on untreated, stressed, and one or two-hour recovered cells (**Methods, Supp. Figure 3E**). As expected, RNAs are generally translationally repressed upon sodium arsenite treatment (**Supp. Figure 3E, F**), with the strongest repression occurring an hour post-treatment, and protein production recovering partially 1-2 hours after stress removal (**Supp. Figure 3G**). Importantly, we observed that stress granule-enriched RNAs show higher SHAPE reactivities (more single-stranded) than stress granule-depleted RNAs (**Figure 2C**) upon stress, agreeing with the results of our structure probing inside stress granules. Inversely, we also observed that RNAs that are more single-stranded (high accessibility) are more enriched in stress granules as compared to RNAs that are more double-stranded (low accessibility) (**Figure 2D**). To examine the RNA structures of stress granule-enriched and -depleted RNAs carefully, we plotted the reactivities of a representative long-noncoding RNA (NORAD), that has been shown to enrich in stress granules (Matheny et al., 2021). NORAD RNA indeed becomes more single-stranded upon sodium arsenite stress (**Figure 2E, Supp. Figure 4A**). In contrast, a stress granule-depleted mRNA, GAPDH (Khong et al., 2017), becomes more double-stranded upon stress (**Figure 2F, Supp. Figure 4B**). Intriguingly, across the four time points, stress granule-enriched RNAs are less double-stranded and largely recovered their original structures within two hours of recovery, whereas stress granule-depleted RNAs only recover partially (**Figure 2G**), suggesting that temporal storage of RNA in stress granules can facilitate quick cellular recovery from acute stress.

With the information that stress granule-enriched RNAs are generally more single-stranded, this begs the question of whether stress granule-enriched RNAs tend to be more single-stranded to begin with and/or they become more single-stranded inside the stress granules due to the unwinding of stress granule-resident RNAs. In the following sections, we address these two mechanisms and show that both mechanisms are involved in explaining why stress granule-resident RNAs are more single-stranded than stress granule-excluded RNAs.

### Single-stranded RNAs are preferentially enriched in stress granules

To determine whether stress granule-enriched RNAs are more single-stranded in prior to stress, we calculated the icSHAPE reactivities of stress granule-enriched and -depleted RNAs in unstressed cells. Interestingly, we observed that the stress granule-enriched RNAs are more single-stranded than stress granule-depleted RNAs even before stress (**Figure 2H**), and this increased reactivity is associated with the 3’UTRs and CDS regions of stress granule-enriched RNAs (**Figure 2I**).

To test whether the single-strandedness of an RNA indeed confers better stress granule enrichment, we transfected two RNAs of identical length and GC composition, but with different structuredness into Hela cells (**Figure 2J**). We then stressed the cells and performed stress granule fractionation followed by qRT-PCR analysis to determine their relative abundance inside stress granules versus inside the soluble cytosol. Interestingly, we observed that a higher proportion of the single-stranded RNA is present inside stress granules as compared to the transfected double-stranded RNA (**Figure 2K**), confirming that single-strandedness facilitates better stress granule enrichment during stress.

### The stress granule environment promotes single-strandedness of RNAs

To test the role of stress granule environment in affecting RNA structuredness, we induced stress granules and performed structure probing on stress granule-deficient and stress granule-competent U-2 OS cells **(Figure 3A)**. To generate stress granule-deficient U-2 OS cells, we engineered cells to express a peptide that encodes the first 40 amino acids of USP10, which is known to block stress granules by reducing G3BP1/2 valency (Kedersha et al., 2016; Sanders et al., 2020) with a tunable degron (Banaszynski et al., 2006). It is worth noting that the key advantage of cells expressing the USP10 peptide over G3BP1/2 dKO cells is that it increases specificity toward studying SG biology and not other SG-unrelated G3BP functions. As expected, adding shield-1 increases USP10 peptide expression and blocks stress granule formation (**Figure 3B, Figure Supp. Figure 5A, 5B**). The results confirmed the successful establishment of the cell line system that can be used for probing RNA structures in the presence and absence of shield-1, thereby comparing the role of stress granules in the modulation of RNA structure (**Figure 3A**).

**Figure 3.**
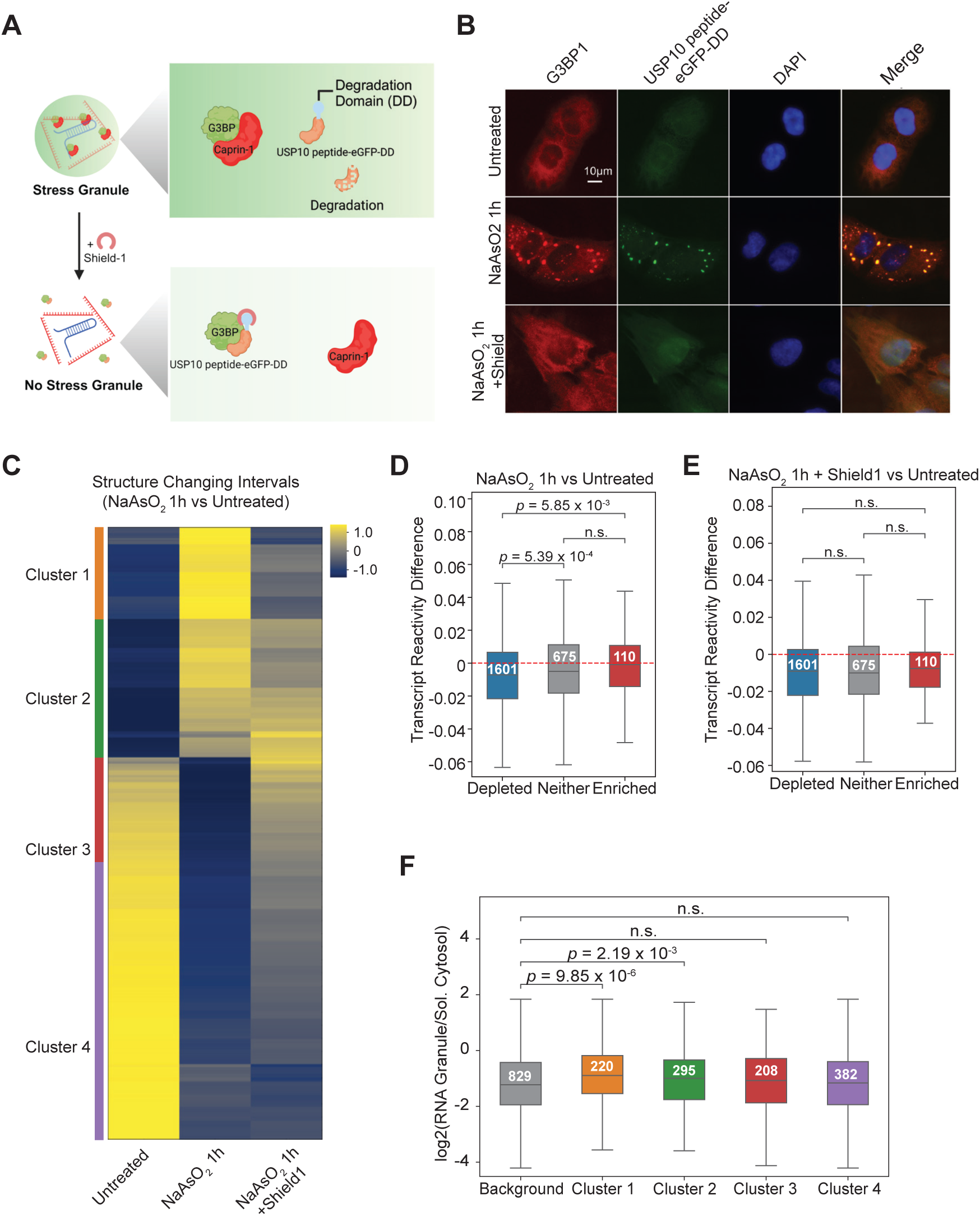
RNA single-strandedness after sodium arsenite treatment is dependent on stress granule formation. **A**, Schematic of the USP10-peptide-degradation system in U-2 OS cells. **B**, Immuno-fluorescence staining of G3BP1 (red), USP10-GFP (green) and nuclei (labeled as DAPI in blue) upon sodium arsenite-induced stress in the presence or absence of Shield-1 (500nM) for 24h. **C**, Heatmap showing reactivity of RNA regions in unstressed U-2 OS cells, and in cells treated with sodium arsenite with and without Shield-1 (500nM). **D,E**, Boxplot showing the distribution of changes in reactivity in stress-granule enriched, depleted and neither transcripts before and after sodium arsenite treatment (**D**), as well as with and without Shield-1 after sodium arsenite treatment (**E**) in U-2 OS cells. **F**, Boxplots showing the distribution of RNA stress-granule-enrichment in Clusters 1-4 in (**C**). Statistical analysis of **D-F** was done with Mann-Whitney U Test with sample sizes and p-values labelled in the panels.

Our quality check against the known structure of 28S rRNA reflected a good accuracy (AUC-ROC > 0.7, **Supp. Figure 5C**). As expected, sodium arsenite treatment of U-2 OS cells resulted in a transcriptome-wide increase in structuredness, similar to what we observed above for HeLa cells (**Figure 3C**). Meanwhile, we also observed that stress granule-enriched RNAs are resistant to the trend of increasing structuredness upon stress, as compared to the stress granule-depleted RNAs in U-2 OS cells (**Figure 3D**). However, stressing the cells and yet not allowing the stress granules to form by adding shield-1 resulted in a reversal of single-strandedness in transcripts that are enriched in stress granules (**Figure 3E**). These results suggest that stress granules are involved in limiting the folding of stress granule-resident RNAs that occur for a wide spectrum of transcripts during stress.

To get more insights into the underlying role of SG in the modulation of RNA structure, we clustered RNA regions according to their reactivities. We observed a cluster, named as Cluster 1, became single-stranded upon stress in the presence of stress granules, and yet remained double-stranded in the absence of stress granule (shield-1 treatment, **Figure 3C, 3F**). Interestingly, the RNAs in this Cluster1 were the topmost enriched in stress granules among others (**Figure 3F**), further supporting the observation that the stress granule environment may facilitate single-strandedness along RNAs.

### Stress granule-enriched RNAs form fewer intra- and inter-molecular RNA interactions

To explore the relatively reduced RNA structure in stress granule resident RNAs, we asked if the reduced RNA structure is due to fewer intra-molecular RNA interactions and/or inter-molecular RNA interaction via RNA proximity ligation experiments (SPLASH) (Aw et al., 2016). We performed two biological replicates of SPLASH on untreated and arsenite-treated cells (**Supp. Figure 6A**), and obtained intra- and intermolecular pair-wise RNA-RNA pairing information for 1,497 and 3,105 genes in control and stressed conditions, respectively (**Supp. Figure 6B**). We observed a good correlation between biological replicates (**Supp. Figure 6C**), and a high AUC-ROC based on the secondary structure of 28S rRNA (**Supp. Figure 6D**), indicating that our SPLASH data is of good quality.

Similar to what we observed in icSHAPE, SPLASH analysis revealed that pair-wise RNA-RNA interactions are increased during stress for both intra- and intermolecular RNA-RNA interactions **(Figure 4A, B)**. In addition, we found that stress granule-enriched RNAs are less paired in both intra- and inter-molecular interactions than depleted RNAs **(Figure 4C, D)**. We also observed that the stress granule-enriched RNAs have shorter intramolecular interactions normalized to transcript length, as compared to depleted RNAs **(Figure 4E)**. Shorter local interactions tend to recover quickly after unwinding, and hence tend to be more stable than longer interactions, suggesting that stress-granule-enriched RNAs could also be actively unwound inside stress granules.

**Figure 4.**
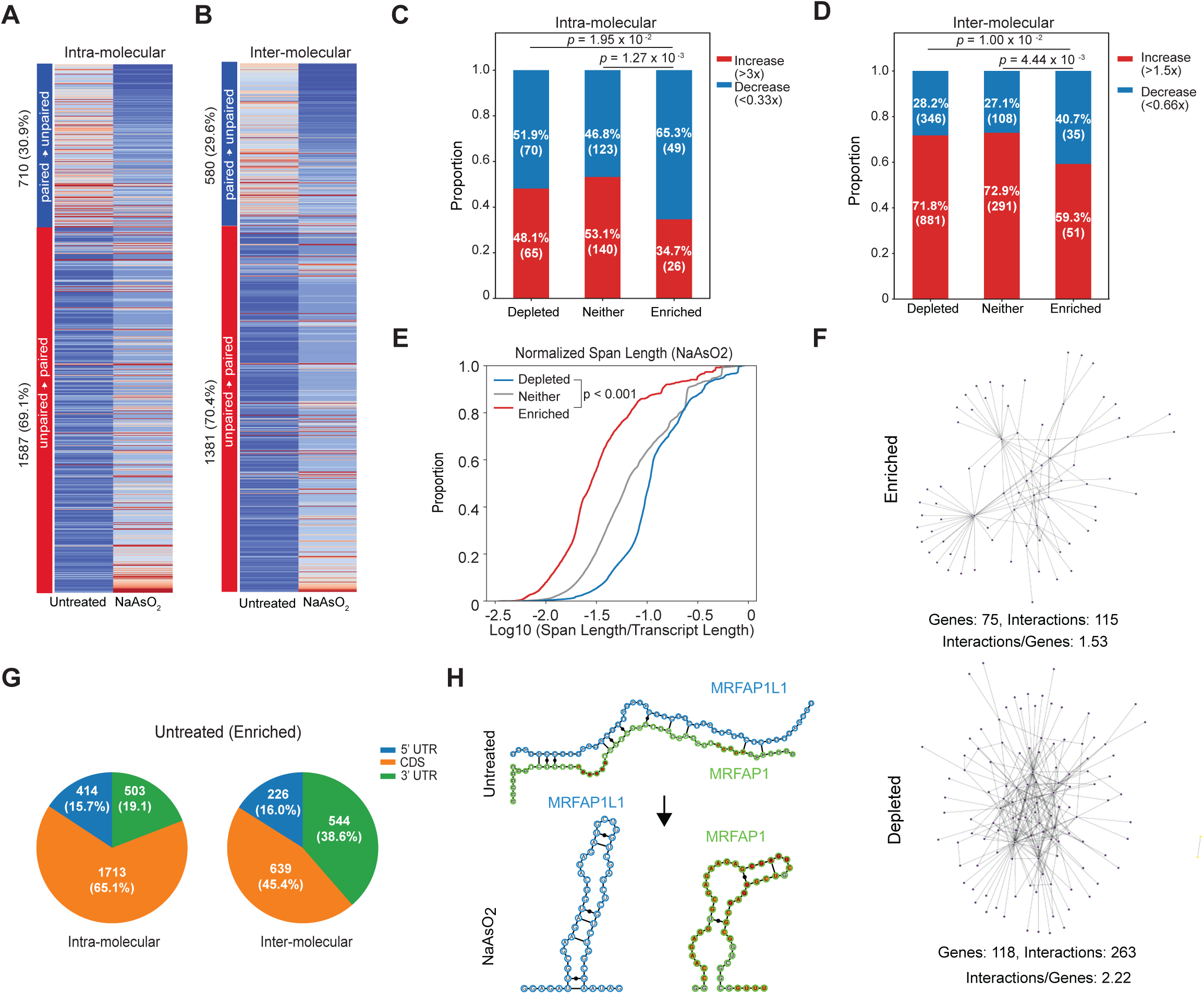
Stress granule-enriched RNAs have fewer number of intermolecular RNA-RNA interactions. **A,B**, Heatmaps showing the number of SPLASH intramolecular interactions (**A**) and intermolecular interactions (**B**) in unstressed and sodium arsenite stressed Hela cells. **C, D**, Stacked barplots showing the percentage of intramolecular (**C**) and intermolecular interactions. Statistical analysis of **C,D** was done using Chi-square Test, with p-values labelled in the panels. (**D**) that changed upon sodium arsenite treatment. **E**, Cumulative plots showing the proportion of intramolecular chimeric interactions that span a certain distance along stress granule enriched (red), depleted (blue), and no preferential localization (grey) mRNAs. Statistical analysis of **E** was done using Mann-Whitney U Test, with p-value labelled in the panel. **F**, RNA-RNA interaction network of stress granule-enriched (left) and depleted (right) RNAs in unstressed Hela cells. **G**, Pie charts showing the distribution of intramolecular and intermolecular interactions that fall into 5’-UTR, CDS regions, and 3-’UTRs of stress-granule-enriched transcripts. Statistical analysis of **G** was done using Chi-square Test, with p-values labelled in the panel. **H**, Structural models of an inter-molecular RNA-RNA interaction between MRFAP1 and MRFAPL1 as well as their local structures using RNAfold. Higher icSHAPE reactivity is shown as red.

To observe how the RNAs interact with each other in the form of intermolecular RNA-RNA interaction networks in untreated and stressed cells, we plotted RNA-RNA interactions between different RNAs for stress granule-enriched and -depleted RNAs. Interestingly, we observed that stress granule-enriched RNAs form fewer intermolecular RNA interactions as compared to depleted RNAs, even in the unstressed cells (**Figure 4F, Supp. Figure 6E**), suggesting that stress granule-enriched RNAs are naturally less paired intermolecularly. To determine the distribution of pair-wise RNA-RNA interactions along the mRNAs, we plotted the location of the pairings along the mRNAs. Interestingly, we observed that stress granule enriched RNAs form more intramolecular RNA-RNA interactions in the coding region and more intermolecular RNA-RNA interactions using their 3’UTRs before stress (**Figure 4G**), and that these features are maintained after stress induction (**Supp. Figure 6F**).

To show how intermolecular RNA-RNA interactions can be remodeled, we examined an intermolecular RNA-RNA interaction between two stress granule-enriched RNAs MRFAP1 and MRFAP1L1. MRFAP1 and MRFAP1L1 RNAs showed strong RNA-RNA base pairing evaluated by SPLASH under control conditions and a 1.5-fold decrease in interactions during stress (**Figure 4H**). Co-fold analysis constrained by icSHAPE reactivity showed that MRFAP1 can form extensive RNA-RNA interactions with MRPAP1L, suggesting these two RNAs pair well with each other under unstressed conditions (**Figure 4H**). As the respective RNA sequences from MRFAP1 and MRFAP1L1 tend to form strong local structures in the presence of stress (**Figure 4H**), we hypothesize that the disruption of intermolecular pairing inside the stress granules likely shifts the pairing equilibrium towards more local structures.

### SRSF1 binds to single-stranded regions along RNA to influence stress granule enrichment

As RNA binding proteins (RBPs) are important cellular factors that can regulate RNA structures, we tested for the involvement of RBPs in regulating RNA structures during stress by performing motif enrichments and also checking for enrichments using eCLIP data that are publicly available. To determine whether any of the RBPs are involved in the structure remodeling of stress granule-associated RNAs, we first focused on candidate RBPs that could alter RNA structures in SGs (**Figure 5A, B**). We identified 33 RBPs that are enriched for our structure-changing regions from the eCLIP data, as well as 196 motifs in 67 RBPs using motif enrichment (**Figure 5A, B**). Overlapping the two enrichment analyses resulted in three RBPs (SRSF1, SRSF9, and FXR2), indicating the binding of these RBPs is associated with the stress-induced RNA structure changes (**Figure 5A, B**).

**Figure 5.**
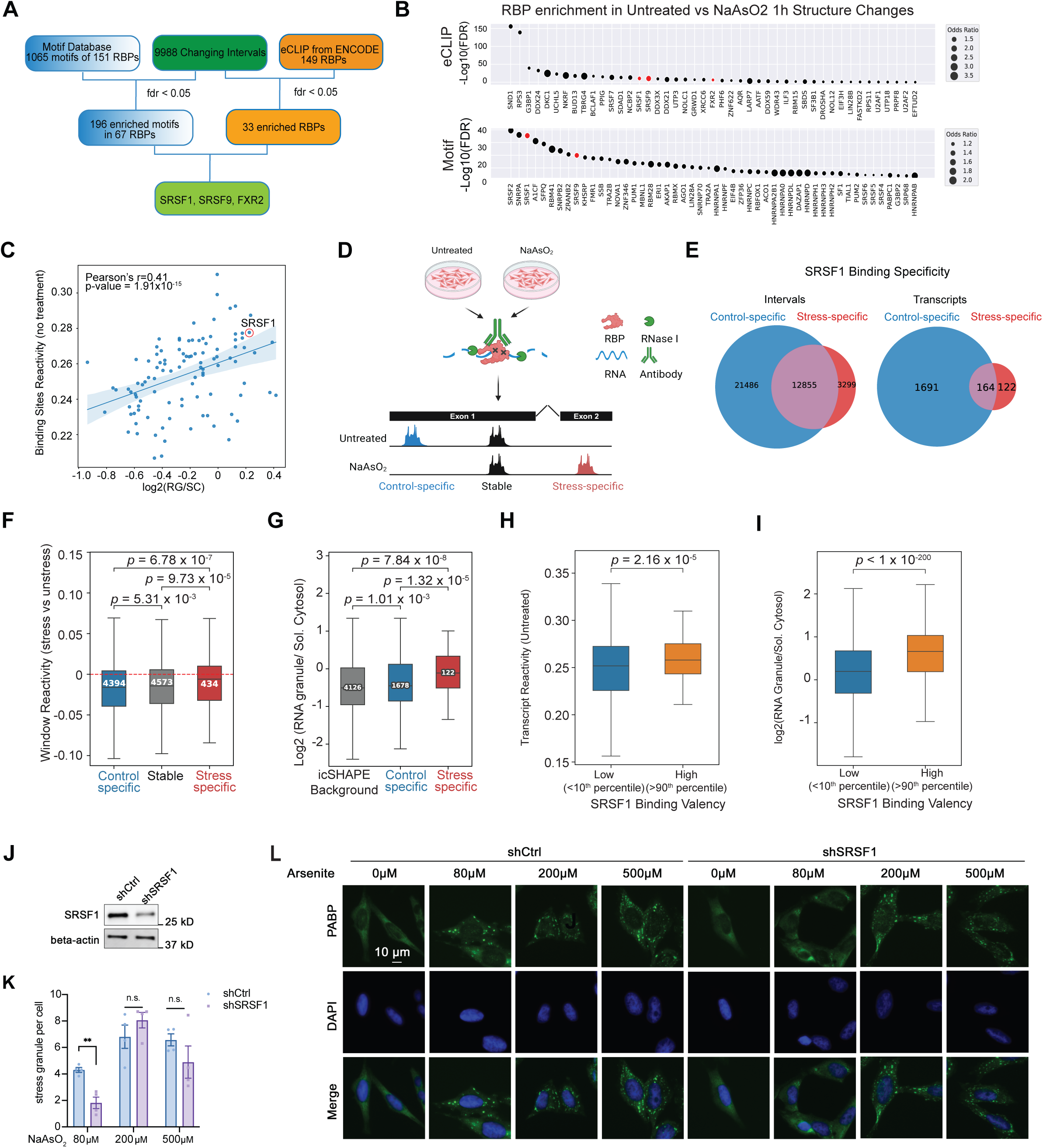
SRSF1 binds to single-stranded regions along RNAs for stress granule formation. **A,** RBP enrichment analysis of differential structure changing regions during sodium arsenite stress using motif enrichments and eCLIP enrichments. **B**, Dotplots showing the significant RBP candidates obtained from the enrichment analysis using eCLIP database (top) and motif database (bottom), respectively. **C**, Scatterplot showing the correlation between binding site reactivity (single-strandedness) and stress granule enrichment of RBP target transcripts. SRSF1 target transcripts were circled in red. **D**, Experimental workflow of using eCLIP to identify SRSF1-binding sites in unstressed or stressed Hela cells. **E**, Venn diagram showing the overlap of RBP-binding peaks identified in unstressed and arsenite-stressed cells. SRSF1 binding sites that are present only in unstressed cells are called “control specific”, only in stressed cells are called “stress specific” and in both unstressed and stressed cells are called “stable”. **F**, Boxplot showing the distribution of reactivity differences in control specific, stress specific, and stable SRSF1 binding regions, before and after sodium arsenite treatment. **G**, Boxplot showing the distribution of stress granule enrichment in SRSF1 target RNAs that are only bound to SRSF1 in unstressed cells or SRSF1 in stressed cells. **H**, Boxplot showing the distribution of transcript reactivities in SFSR1 targets with high and low number of SRSF1 binding sites (cutoff = 90%). **I**, Boxplot showing the distribution of stress granule enrichment for SRSF1 targets with high and low number of SRSF1 binding sites (cutoff = 90%). Statistical analysis of **F-I** was done using Mann-Whitney U Test, with p-values labelled in the panels. **J,** Western blot analysis of SRSF1 knockdown. **K,L**, Immunofluorescence staining (**K**) and quantification of stress granule numbers (**L**) in HeLa cells after treating the cells with different concentrations of sodium arsenite, with and without SRSF1 knockdown. PABP (indicating stress granules) was labeled in green. Nucleus were stained with DAPI in blue. Statistical analysis of **L** was performed using the Student’s T-Test. Data are shown as mean ± SEM, n = 5. *p < 0.05, **p < 0.01, ***p < 0.001.

To our surprise, two of the SR proteins (serine/arginine-rich proteins), namely SRSF1 and SRSF9, were shortlisted. Although SRSF1 and SRSF9 predominantly localize in the nucleus to function in RNA splicing, they could also shuttle between the cytoplasm and nucleus and have functions beyond splicing (Sanford et al., 2004). While in the cytoplasm, SRSF1 interacts with stress granule nucleator protein, TIA1, to modulate mRNA translation, and localize to stress granule upon stress (Delestienne et al., 2010; Twyffels et al., 2011). To validate the stress granule localization of SRSF1/9 in Hela cells, we performed immunofluorescence staining in HeLa cells that stably express EGFP-G3BP fusion protein. While it was notable that SRSF1 and SRSF9 predominantly in the nucleus, they co-localize with G3BP1 in the stress granules upon arsenite stress (**Supp. Figure 7A, B**).

We then sought to understand the association between RBP binding, RNA structure, and RNA stress granule enrichment. By plotting the single-stranded propensities of RBP targets against their enrichment in the stress granules, we observed a positive correlation between the reactivity of the RBP binding site and the SG-enrichment. This supports our prior observations that single-stranded RNAs tend to be more enriched in stress granules (**Figure 5C**). Interestingly, we observed that the targets of one of our enriched RBPs, SRSF1, also tend to be more single-stranded among the RBPs (**Figure 5C**), indicating a mechanism that SRSF1 could bind to single-stranded RNAs to recruit them to the stress granules.

To profile the locations of SRSF1-binding in the transcriptome, we performed two biological replicates of eCLIP experiments in untreated and stressed Hela cells (**Figure 5D**). We observed that the reads from the two biological replicates are highly correlated (**Supp. Figure 8A**). Motif enrichment analysis of the enriched peaks suggested that SRSF1 binding peaks are enriched for the known SRSF1 motif (**Supp. Figure 8B**), and that SRSF1 predominantly binds to the coding region in both unstressed and stressed conditions as expected (**Supp. Figure 8C**), indicating that our eCLIP data are of good quality. In total, we identified 37,640 peaks that bind to SRSF1 in both untreated and stressed conditions, with 57.1% (21,486 regions in 1,691 genes) of the peaks being present in untreated cells only, and 34.2% (12,855 regions, 164 genes) of the peaks being present in both control and stressed cells and 8.8% (3,299 regions, 122 genes) of the peaks being present in stressed cells only (**Figure 5E**). Metagene analysis showed that true SRSF1 binding regions, namely those with eCLIP evidence, are indeed more single-stranded than regions that contain the same motif sequence, but without eCLIP binding evidence, suggesting that SRSF1 is associated with single-stranded regions in both untreated and stressed cells (**Supp. Figure 8D, E**). Importantly, RNA regions that are bound to SRSF1 during cellular stress showed an even stronger single-stranded propensity than regions that are consistently bound to SRSF1 (**Figure 5F**) and are more enriched in the stress granules (**Figure 5G**). As SRSF1 binding correlates with single-strandedness along a transcript, we tested whether transcripts with more SRSF1 binding sites are less structured. Transcripts were stratified by SRSF1 binding site abundance for analysis. Interestingly, we observed that transcripts with more SRSF1 binding sites are indeed more single-stranded (**Figure 5H**), even when they are controlled for transcript length (**Supp. Figure 8F**). Moreover, more SRSF1 binding sites are associated with greater stress granule enrichment, indicating the novel role of SRSF1 in directing its binding targets to stress granules in an RNA structure-dependent manner. (**Figure 5I, Supp. Figure 8G**).

To determine whether SRSF1 is functionally important for stress granule formation, we stressed the Hela cells and counted the number of stress granules formed in untreated and SRSF1 knockdown cells **(Figure 5J)**. Interestingly, while SRSF1 knockdown does not impact stress granule formation under high-stress conditions (200µM above), we observed that silencing SRSF1 reduces the number of stress granules formed under low-stress conditions (80µM), suggesting that SRSF1 is indeed important for stress granule assembly, albeit in a manner depending on stress intensity (**Figure 5K, L**). To understand the potential role of SRSF1-binding during stress response at a cellular level, we knocked down SRSF1, applied acute SA stress, and examined cell viability two days post-stress. Interestingly, we observed that depletion of SRSF1 led to higher cell viability, although the precise mechanisms of how it does so warrant future investigation (**Supp. Figure 8H**).

### RNA helicases unwind RNA structures inside stress granules

In addition to single-stranded RNAs being preferentially recruited into stress granules, another hypothesis for why RNAs are more single-stranded within SGs is that they are being actively unwound in stress granules. As many RNA helicases are enriched inside the stress granule (Jain et al., 2016), we asked if these RNA helicases are involved in unwinding SG-resident RNAs. We first performed ATP-depletion by treating cells with 2-Deoxy-D-Glucose (2-DG) and carbonyl cyanide 3-chlorophenylhydrazone (CCCP) that inhibit glycolysis and oxidative phosphorylation, respectively, enabling a global inhibition of RNA helicase activity (**Figure 6A**). We then tested the extent of RNA structuredness in the RNA granule before and after ATP depletion (**Figure 6A, Methods**). We confirmed that ATP depletion decreased the amount of ATP inside cells dramatically (**Figure 6B**), and that our icSHAPE libraries are of good quality (**Supp. Figure 9A**). In contrast to the single-stranded propensity of RNAs inside stress granules during stress, inhibiting helicases by ATP depletion results in a similar level of structuredness of RNAs inside stress granules as those in the soluble cytoplasm (**Figure 6C, D**), suggesting that RNA helicases play an important role in unwinding RNA structures inside the stress granules.

**Figure 6.**
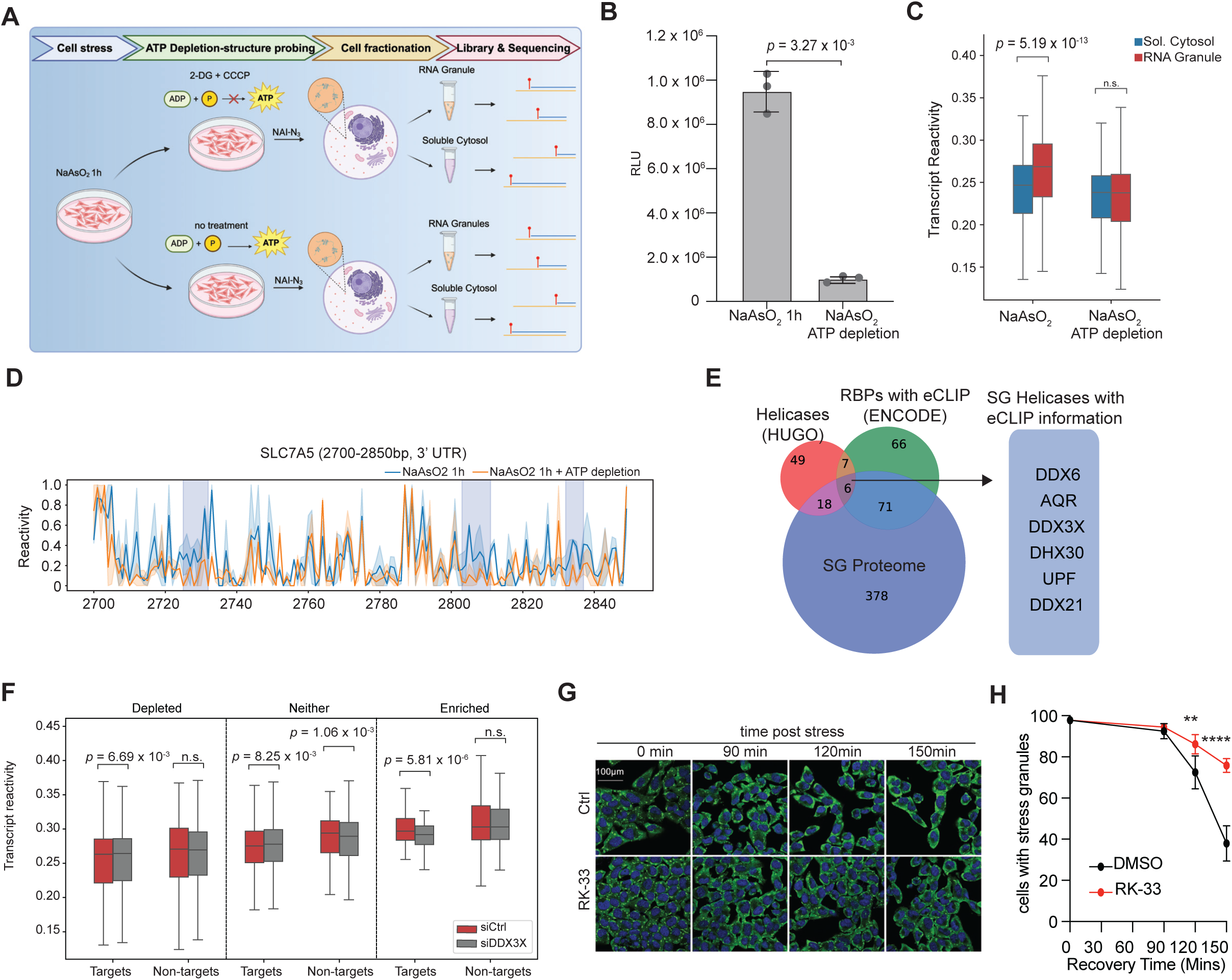
DDX3X unwinds RNAs inside stress granules to enable efficient stress granule disassembly. **A**, Workflow of probing RNA structures inside stress granules after depletion of ATP. Briefly, we depleted ATP after treating the cells with sodium arsenite using 2-DG (10mM) and CCCP (0.1mM). We then performed NAI-N_3_ labeling, subcellular fractionation, and library preparation. **B**, Barplots showing the amount of ATP levels present inside cells with and without ATP depletion. In cell ATP amount is measured by cell-titer Glo. **C**, Boxplots showing the distribution of transcript reactivity in stress-granule RNAs versus cytospolic RNAs, in the presence and absence of ATP depletion. Statistical analysis of **C** was done using Mann-Whitney U Test, with p-values labelled in the panel. **D**, Line plots showing the reactivities of a representative stress granule enriched transcript, in the presence and absence of ATP. Blue intervals represent significant decrease in single-strandedness upon depletion of ATP. **E**, Venn diagram showing the overlap in genes found in three public databases, HUGO helicases (Umate et al., 2011), ENCODE eCLIP RBPs, and stress granule databases (Jain et al., 2016). **F**, Boxplots showing the reactivity of DDX3X target and non-target RNAs that are stress granule-enriched, depleted, or neither, in control and DDX3X knockdown cells. Statistical analysis of **F** was done using Mann-Whitney U Test, with p-values labelled in the panel. **G,H**, Immunofluorescence staining (G) and quantification (H) of stress granules after sodium arsenite treatment, in the presence of absence of DDX3X inhibitor, RK-33 (6µM), at the indicated time points U-2 OS cells expressing mutant USP10 peptide (see materials and methods). Statistical analysis of **B,H** was performed using the Student’s T-Test. Data are shown as mean ± SEM, n = 5. \**p* < 0.05, \*\**p* < 0.01, \*\*\**p* < 0.001, \*\*\*\**p* < 0.0001.

We next sought to identify the helicases involved in unwinding the SG-resident RNAs. Toward this, we first overlapped the list of helicases with those present in the stress granules and with available eCLIP data (**Figure 6E**). This resulted in a total of six helicases, including DDX6, AQR, DDX3X, DHX30, UPF, and DDX21. As DDX3X is involved in the assembly of stress granules (Cui et al., 2020), we tested its role in regulating RNA structures more extensively.

To confirm that DDX3X is indeed important for unwinding RNAs in stress granules, we knocked down DDX3X using siRNA and performed structure probing in untreated and sodium arsenite-stressed cells. As expected, the expression of DDX3X was downregulated upon siRNA knockdown (**Supp. Figure 9B**), and our icSHAPE libraries are of good quality (**Supp. Figure 9C**). Importantly, the downregulation of DDX3X partially disrupts the single-stranded propensity of its targets in stress granule-enriched RNAs without influencing the structure of stress granule-depleted transcripts (**Figure 6F**), indicating that DDX3X is responsible, at least partially, for unwinding RNAs inside stress granules.

Stress granule assembly and disassembly are critical for stress response at the cellular level. As promiscuous RNA-RNA pairing would require costly helicase-mediated unwinding to restore RNA function post stress, we hypothesized that RNAs are actively maintained as relatively single-stranded inside stress granules to enable rapid stress granule disassembly and cellular recovery. As such, inhibition of DDX3X could result in slower cellular recovery in the absence of stress. To test this, we stressed the cells and examined the amount of stress granules remaining after stress removal in the presence and absence of DDX3X inhibition, using a specific compound RK-33 that targets DDX3X. Interestingly, we observed that inhibiting DDX3X resulted in slower stress granule dissolution after stress removal (**Figure 6G,H**), suggesting that increased RNA-RNA pairing in stress granules impairs the rate of stress granule disassembly.

## Discussion

Being able to respond to environmental stress quickly is the key for organismal survival. Here, we have presented using multiple orthogonal evidence that RNAs tend to be less structured inside stress granules as compared to outside of the stress granules during acute arsenite stress. The finding is in line with a previous report that single-stranded RNAs but not double-stranded RNAs undergo LLPS in a manner dependent on G3BP1 (Yang et al., 2020). The unstructured state may have significant consequences for gene expression regulation, as RNA structures are known to influence both translation and decay (Su et al., 2018; Wang et al., 2021). Mechanistically, we provided evidence showing that it is partially attributed to the preferential recruitment of single-stranded RNAs into the stress granules by RBP such as SRSF1. We found that silencing of SRSF1 impaired stress granule formation at low levels of stress, suggesting that it could play an important role in RNA partitioning. Additionally, we observed active unwinding of RNA structures by helicases inside stress granules, and showed that inhibition of a specific helicase DDX3X resulted in increased RNA structures in the stress granules and slower dissolution of stress granules upon removal of stress, slowing down cellular recovery upon stress. Our study highlights the importance of regulating RNA-RNA pairing in response to enable efficient response to cellular stress.

Several models have been proposed to explain the role of RNA in SG condensation. While transcript length and AU richness has been shown to be important for stress granule enrichment (Khong et al., 2017), how RNAs utilize their structures for stress granule formation is largely unknown. One prominent model suggests that SGs form as a biophysical consequence of RNA aggregation (Ripin and Parker, 2022; Van Treeck et al., (2018); Tauber et al., (2020); Parker et al., (2024); Trussina et al., (2024). Specifically, RNA tends to promiscuously interact with other RNAs when not bound to ribosomes. During rapid translation shutdown, the cellular machinery, including helicases, becomes overwhelmed, leading to SG formation via promiscuous non-specific intermolecular RNA-RNA interactions.

Our findings present an intriguing challenge to the RNA aggregation model. While we anticipated an increase in RNA structure and intermolecular RNA-RNA interactions for RNAs localized to stress granules, our icSHAPE and SPLASH data revealed the opposite: stress granule-enriched RNAs are less structured and paired intermolecularly as compared to SG-depleted RNAs. Intriguingly, our data raise the possibility that minimal RNA-RNA interactions are sufficient to promote aggregation of biomolecules and thus stress granule formation, highlighting the essential role of RNA-binding proteins in regulating this process.

Interestingly, our data align with recent studies indicating that RNAs enriched in stress granules are preferentially targeted by ATP-dependent processes, helicases, and RNA-binding proteins (Tauber et al., 2020; Trussina et al., 2024). This could be interpreted as a mechanism to mitigate promiscuous RNA aggregations that makes it difficult to reverse the stress granule formation process during stress.

Loss of DDX3X function, mutations in DDX3X, or overexpression have been linked to various diseases, including neurodevelopmental disorders and cancers **(Cruciat et al., 2013; Samir et al., 2019)**. Additionally, DDX3X-mutant SGs can also disrupt global translation, unlike classical arsenite-induced stress granules **(Valentin-Vega et al., 2016)**. Here, by showing that inhibition of DDX3X results in increased structuredness of its target RNAs in stress granules and slows down cellular recovery after stress removal, we linked RNA structure regulation to potentially different disorders that are associated with DDX3X mutant phenotypes. Consistent with this finding, Trussina and colleagues recently reported that inhibition of DDX3 accelerates SG formation and delays their disassembly (Trussina et al., 2024). We anticipate that more studies will elucidate the relationship between RNA structure and cellular functions in physiological and pathological contexts.

In summary, by providing the first comprehensive RNA structurome and RNA-interaction map for stress granules, we provide significant insights into how RNAs are organized inside stress granules. We believe this will help address the ongoing debates surrounding how RNAs are recruited to and promote SG condensation, as well as how SGs regulate RNA fate. Moreover, we anticipate that further studies will reveal how manipulating RNA structures within SGs can alter SG biophysical properties, gene expression, and disease biology, providing mechanistic insights into the role of stress granules in diverse pathological settings, such as neurodegenerative diseases and cancers.

## Materials and Methods

### Cell culture and stress treatment

HeLa, HEK293T, and U2-OS cells were purchased from ATCC and maintained in high glucose DMEM supplemented with 10% fetal bovine serum, and 1% penicillin/streptomycin at 37°C/5% CO_2_. One day before stress treatment, the medium replenished with fresh DMEM. On the day of treatment, half of the used medium were collected, stored at 37 °C, and used during recovery. Sodium Arsenite (0.5M) were diluted 100 times in fresh DMEM, and then added in the culture medium at the ratio 1:10 to achieve final 500µM final concentration unless otherwise specified. U-2 OS cells stably expressing USP10-peptide-eGFP-DD, were generated using lentiviral transduction of plasmids encoding the first 40 amino acids of USP10 fused to eGFP-DD. Following transduction, cells were sorted via FACS to select for high expressors. The first 40 amino acids of USP10 have been previously demonstrated to inhibit stress granule formation, whereas the mutant variant (Kedersha et al., 2016).

### Immunostaining and Fluorescence microscopy

HeLa or U-2 OS cells were seeded on glass cover slides before cells were stressed and washed with cold phosphate-buffered saline (PBS) followed by fixation with cold 4% paraformaldehyde solution in PBS (santa cruz, sc-281692) for 12min. Cells were then washed with PBS and permeabilized with PBS supplemented with 0.2% Triton-X100 for 15min. The fixed cells were blocked using PBS supplemented with 1% BSA, then incubated with following antibodies at 4°C overnight: PABP (1:60; Santa cruz, sc-32318); G3BP1 (1:300; Cell Signaling Transduction #17798S). Fixed cells were incubated with secondary antibodies (1: 1,000; Alexa 488/594, Life Technologies) and DAPI (NucBlue, Invitrogen, R37606; 1 drop per 500 µL) at room temperature for two hours. After three washes with PBST, 200 µL PBS was added and the slides were observed under fluorescent microscope. Stress granules were counted using Image J using “analyze particle” function with the restriction of following parameters: size = 4 to 100-pixel, circularity = 0.8-1.0.

### Stress granule recovery experiment

U-2 OS cells were subjected to 500 µM sodium arsenite treatment for 1 hour. Following the stress treatment, cells were washed and maintained in either 6 µM RK-33 or DMSO as a control. At designated timepoints, cells were fixed and stained with the stress granule marker G3BP1 (ab56574, Abcam). Images were acquired using a confocal microscope. To ensure unbiased analysis, sample images were blinded prior to manually counting cells containing stress granules.

### Plasmid construction and siRNA knock down

For shRNA constructs, shSRSF1 sense and shSRSF1 antisense were annealed ligated into the pLKO.1-puro plasmid. For synthetic dsRNA and ssRNA constructs, the DNA templates were synthesized using gblock method (IDT). The synthesized DNA fragments were subjected to Gibson assembly (NEB, E2611L) to clone into a pmiR-Glo plasmid. To generate RNA fragments for transfection, plasmids were linearized, in vitro transcribed (NEB, E2040S) and gel purified (NEB, T2050S) before quantitation using nanodrop. For siRNA knockdown, a pool of three siRNAs against DDX3X (s4004, s4005, s4006) were purchased from Ambion.

### RNA structure probing (icSHAPE)

Cells were cultured in 15 cm dish till 80% confluency and treated with arsenite as described above. After stress, cells were washed once with 10ml PBS, trypsinized and washed with another 5ml PBS. Cells were pelleted by centrifugation at 300g for 5 min. PBS was aspirated, the pellets were resuspended in PBS supplemented with 20mM NAI-N_3_ or 1% DMSO and transferred to a 1.7ml tube, which was followed by rotating the cell suspension for 5min at 37°C end-to-end. Cells were pelleted at 300g for 3min. The cell pellet was lysed in Trizol and total RNA was isolated following the standard trizol protocol. icSHAPE library preparation was carried out according to the protocol described before with minor modifications.

### RNA structure analysis in subcellular components (Frac-SHAPE)

Mammalian RNA granule was enriched using the methods adapted from the protocol described previously (Khong et al., 2016; Namkoong et al., 2017). Cells were treated as described above and were subjected to differential centrifugation to separate RNA granule (RG) and soluble cytoplasm. Briefly, after the DMSO/NAI-N_3_ treatment, cells were washed with PBS and pelleted at 300g for 3min. Cells were then lysed in 1ml lysis buffer L (50 mM Tris pH 7.6, 50 mM NaCl, 5 mM MgCl_2_, 0.2% NP-40) by passing through the 25 5/8G needle 4-5 times. Lysate were centrifuged at 1000 × g for 5 minutes and the supernatant were immediately transferred to (∼900µL, cytoplasmic extract) to a clean pre-chilled tube on ice. The supernatant were further centrifuged at 18,000g for 20min to separate RNA granules from the soluble cytoplasmic fraction. The RG fraction was washed with 300ul buffer L and resuspended 150ul buffer L, after which, a final centrifugation at 1,000g for 2min was applied to remove debris. RNA isolation were performed by isolated by adding 3 vol of Trizol LS to each fractions prepared, which is followed by the RNA isolation using Trizol and RNeasy purification.

### icSHAPE analysis in vitro and in vivo

Cells were subjected to DMSO/NAI-N_3_ treatment followed by icSHAPE library preparation described before (Flynn, et al, 2016). For in vivo SHAPE experiment, total RNA were extracted using Trizol method. Upon mRNA selection, RNA were denatured at 90°C, quickly put on ice for 2min, add 5xSHAPE structure probing buffer (250mM Tris-HCl pH7.4, 50mM MgCl2, 750 mM NaCl), and ramp up from 4°C to 37°C at 0.1°C/s. All icSHAPE libraries were processed using publicly available software icSHAPE-pipe (version 2.0.0) (Li et al, 2019). Briefly, raw single-end sequencing reads were de-multiplexed using in-house index sequences. Reads were collapsed to remove PCR duplicates using readcollapse. Subsequently, 5’ and 3’ adapters were removed using trim (TrimmomaticSE, version 0.38). Reads were then mapped to human reference genome GRCh38 primary assembly from EMSEMBL (release 98) using mapGenome (STAR, version 2.7.9a). Genomic coordinates of uniquely mapped reads were used to quantify SHAPE reactivity per base using calcSHAPE, based on the sliding window strategy described by Li et al, with default window size (200nt) and step (5nt). Genomic coordinates were converted to transcript coordinates using genSHAPEToTransSHAPE whereby nucleotides with base density higher than 100 were considered as valid. Lastly, transcripts that pass the default filters (averaged RT stop count >2, FPKM >5, number of valid nucleotides >10, ratio of valid nucleotides > 0.1) are used for downstream analysis. Differential Structure Intervals (DSI) were identified using algorithms adapted from DiffScan (Yu et al, 2022). To better understand and fine-tune the software in Python, we developed a Python version of DiffScan, adapted from the original version which is written in R. Briefly, a sliding window (default window size is 5nt) approach was used to calculate the significance for each nucleotide. Nucleotides with absolute reactivity difference >0.2 and p-value <0.05 were considered as significantly different nucleotides. Consecutive differential structure nucleotides that are less than 5nt apart will be merged to form an interval. Finally, intervals with an average reactivity difference of 0.1 and above are considered as differential structure intervals. Intervals with more than 50% reactivity recovery efficiency in either Recover 1h or Recover 2h time points will be considered as recovered DSI.

### Western blot analysis

Cells were cultured in 6-well plates. On the day of harvesting, cells were washed with PBS before being lysed in RIPA buffer (Thermo cat. 89901), supplemented with 1x Protease inhibitor (Roche cat.61711697498001) and 1x Phospho-STOP (Roche cat. 04906845001). Total protein was separated by SDS-polyacrylamide gel electrophoresis and then transferred onto polyvinylidene difluoridemembrane using a wet transfer method (constant current for 2 hours). The membrane was blocked with Intercept® Blocking Buffer in (LI-COR P/N 927-600001) for 1 hour at room temperature, before being incubated with primary antibody at 4°C overnight. Antibodies: Phospho-eIF2α (Ser51) Antibody (1: 1000; CST 9721s), GAPDH (1: 5,000; Protein Tech 60004-1-lg), beta-actin (1: 5,000; Santa Cruz sc-47778), eIF2alpha (1:2000, Bethly A300-712A). The membranes were then incubated with IRDye 680RD (1: 5,000) or 627IRDye 800CW (1:5,000) (LI-COR) secondary antibodies at room temperature for 1 hour before imaging using the LICOR ODYSSEY scanner.

### RT-qPCR analysis

Typically, 1 μg RNA was reverse transcribed using the SuperScript™ IV VILO™ Master Mix (Invitrogen, 11756050) according to the manufacturer’s instructions. The resultant cDNA was diluted by a factor of 10 before 1μL was used for qPCR analysis. qPCR was performed using the ABI-7300 instrument (ThermoFisher Scientific) using the KAPA SYBR FAST qPCR kit (KAPA cat. KK4601) and the gene-specific primers in a final volume of 10 μL. Relative gene expression was measured using the 2^−ΔΔCT^ method and normalized against internal control. The primers used for RT-qPCR can be found below.

### RNA-seq analysis

Gene expression of Untreated and NaAsO_2_ samples was obtained using the DMSO treated samples from icSHAPE experiment. Briefly, raw reads were trimmed using icSHAPE-pipe trim function to remove adapter sequences at 5’ and 3’ ends. Subsequently, trimmed reads were mapped to human reference genome GRCh38 primary assembly with default parameters using STAR (version 2.7.9a). Gene level read counts were obtained using FeatureCounts (version 2.0.1). Differentially expressed genes (DEGs) were identified using DESeq2 (version 1.34.0) with default parameters. Gene Ontology of DEGs was done using Clusterprofiler (version 4.4.2) in R (version 4.1.3) with default parameters. For fractionation samples, DMSO samples of frac-SHAPE were used to quantify RNA Granule enrichment for each gene. Gene-level read counts were obtained by mapping trimmed reads to genome with STAR (default parameters), followed by FeatureCounts. DESeq2 was used to identify genes that are differentially expressed in RNA Granules and Soluble Cytosol fractions. Genes with RNA Granules/Soluble Cytosol foldchange greater than 1.5-fold and adjusted p-value <0.01 were considered as “enriched” whereas genes with RNA Granules/Soluble Cytosol foldchange smaller than 0.66-fold and adjusted p-value <0.01 were considered as “depleted”. The remaining genes are classified as “neither”.

### Structure Modelling of ssRNA and dsRNA

Structure models of ssRNA and dsRNA were obtained using RNAStructure (version 6.4). The function partition was used to calculate the pairing probability. MaxExpect was then used to output structure that maximizes the pairing probability.

### Ribosome profiling library preparation and analysis

Raw reads were trimmed and mapped to genome. Read counts assigned to the CDS region of all protein coding genes were obtained by FeatureCounts. In each of the timepoint (Untreated, NaAsO2 1h, Recover 1h and Recover 2h), Transcript per Million (TPM) of all genes were calculated for ribosome-protected fragments (RPF) and RNAseq background. Translation efficiency is defined by the average RPF/RNAseq from 3 replicates.

### RNA-RNA interaction analysis using SPLASH

SPLASH data pre-processing was done following the computational methods described in Aw et al (2016). Briefly, raw paired-end reads (PE150) were adapter-trimmed and merged using SeqPrep (version 1.3.2). Mergedd reads were then mapped with BWA-MEM (0.7.17) to a curated transcriptome (ENSEMBL release 98) containing the longest isoform whose biotype matches its gene’s biotype. Ribosomal RNA (rRNA) sequences obtained from Protein Data Bank (Acession number: 4V6X) were also included to capture rRNA-related chimeras. Gapped alignments with mapping quality higher than 20 and are at least 10nt apart were kept. Upon mapping the remaining chimeric reads to the genome with STAR (version 2.7.9a), chimeric reads that span an annotated splicing junction were removed. Resultant chimeric reads were binned into pairwise 50nt windows (interactions). Interactions that are present in at least 2 out of the 4 replicates and are with average coverage 1 were kept. The coverage for each interaction was normalized using the normalization module of DESeq2. Finally, the coverage of an interaction is the average of normalized coverage in all replicates. For Inter-molecular Network AnalysisSPLASH Inter-molecular network analysis was done at the gene level. Briefly, inter-molecular chimeras (mapping quality >20) between two genes were summed to obtain gene-level coverage. To remove confounding effect of gene expression, we looked only into highly expressed genes (both genes TPM >200) for downstream analysis. Gene pairs were annotated with RG-localization and classified into “depleted” and “enriched” subgroups. The inter-molecular interactions of the “enriched” and “depleted” subgroups were visualized as an unweighted network using the software python-igraph (version 0.9.6).

### Co-fold Analysis

SPLASH Inter-molecular interactions with high coverage and significant changes upon NaAsO_2_ treatment were shortlisted for further investigation. To validate the robustness of the interactions, the interacting regions were input into RNAcofold (ViennaRNA 2.6.4) to observe their pairing efficiency. icSHAPE reactivities of interacting regions were input into RNAcofold as treatment-specific constraints with parameters m=1.8 and b=0.

### RBP Enrichment of Structure Changing Regions

The curated eCLIP profiles of 150 RBPs were used to overlap with differential structure intervals between NaAsO_2_ and Untreated. The enrichment of RBP binding in structure changing and non-changing regions were tested using Fisher’s exact test. The p-values of all the tested RBPs were corrected using Bonferroni method. RBPs with adjusted p-value <0.05 were considered as enriched for structure changes. Independently, we performed RBP binding enrichment against differential structure intervals using known motifs provided in ATtRACT, a curated motif database. Briefly, occurrences of each motif in the icSHAPE detected transcripts were identified using FIMO from MEME Suite (version 5.4.1) with a cutoff of adjusted p-value < 0.001. Similar to eCLIP RBP binding, the enrichment of motif occurences against differential structure intervals were tested using Fisher’s exact test followed by Bonferroni correction (adjust p-value cutoff = 0.05). Lastly, we identified 3 RBPs that are consistently enriched for structure changing regions by using both eCLIP binding site and motif sequences enrichment approaches.

### eCLIP Data Analysis

Publicly available eCLIP profiles of 150 RBPs in HepG2 and K562 cell lines were downloaded from ENCODE database (accession: ENCSR456FVU) (Van Nostrand et al, 2020). Liftover was performed using *ucsc-liftover* (version 447) to convert hg19 to hg38 coordinates. Genomic coordinates were further converted to transcript coordinates with an in-house script using GTF file from ENSEMBL (release 98). Peaks that span the exons fully and pass the filter of p-value <0.01 and log2(CLIP/Input)>2 were kept. Two replicates from each RBPs were merged using *bedtools merge* (version 2.30.0) with default parameters to obtain a union set of peaks for downstream analysis. In-house eCLIP data was processed using publicly available software i.e. Skipper (Boyle et al., 2023). Briefly, raw sequencing reads were mapped to reference genome downloaded from ENCODE database as suggested by Skipper pipeline (accession: ENCSR425FOI). A partition specific to HeLa expressed genes (TPM >1) was used to generate windows that were to be tested. Windows that pass the default FDR cut-off (FDR<0.2) were considered as enriched for RBP binding. The genomic coordinates were converted to transcript coordinates with an in-house script. A union set of enriched windows of 2 replicates were used for downstream analysis.

## Acknowledgements

We thank all members of Wan Lab for discussion. We also thank Jiaxu Wang for advice, Ziyi Yang for experiment optimization, Natalie Lim for assist in SPLASH library preparation. Y.W. is supported by funding from the A*STAR Investigatorship, the EMBO Young Investigatorship, the National Research Foundation, and the CIFAR Azrieli global scholar fellowship. B.Z. is supported by National Medical Research Council (NMRC) (MOH-OFYIRG23jan-0023).

## Author Contributions

Y.W. conceived and supervised the project; Y.W., A.K. and B.Z. designed the experiments; B.Z. performed cell experiments and J.Z. carried out DDX3X inhibition assay. B.Z. performed library preparation for RNA-seq, Ribo-seq, icSHAPE, SPLASH, and Frac-SHAPE; B.Z. and J.S.X. performed eCLIP library preparation; J.L., Y.Z., Y.W., B.Z., and J.S.X. analyzed the sequencing data; J.L. and R.H. performed RNA structure modeling. B.Z., J.L., A.K., and Y.W. interpreted the data and wrote the manuscript.

## Ethics Statement

The authors declare that this study does not contains any human data.

## Supplemental Figure Legends

**Supplementary Figure 1. Investigation of RNA structure and RNA expression change during sodium arsenite-induced stress. A**, Scatter plots showing the correlation of RT stop sites between biological replicates of indicated treatments. **B**, AUC-ROC plots were obtained by comparing the icSHAPE reactivity to the known accessibility of 28S rRNA. **C**, Volcano plot showing deferentially expressed genes between unstressed and sodium arsenite-stressed Hela cells. **D**, Gene ontology analysis of deferentially expressed genes between unstressed and sodium arsenite-stressed Hela cells. **E**, Pie charts showing the proportion of nucleotides (left panel) and transcripts (right panel) that undergo changes in reactivity upon sodium arsenite stress.

**Supplementary Figure 2. *In vitro* RNA structure probing before and after arsenite-induced stress. A**, Scatter plots showing the correlation of RT stop sites between biological replicates of indicated treatments obtained by *in vitro* SHAPE analysis. **B**, Heatmap showing the reactivity of RNA windows before and after stress when the RNAs are structure probed in vivo (left two columns) and in vitro (right two columns). **C**, Density plots showing the distribution of RNA regions that become more double-stranded or more single-stranded upon sodium arsenite treatment.

**Supplementary Figure 3. RNA structures are more single-stranded inside stress granules than those inside soluble cytosol. A**, Barplots showing the relative enrichment of representative genes that exhibit preferential localization during stress. Briefly, stress granule depleted RNA (UQCRC), enriched RNA (AHNAK) and RNA with no preferential localization (PH4B) were analyzed by RT-qPCR after subcellular fractionation. **B**, Scatter plot showing the correlation of stress granule enrichment between our data and a published dataset. Pearson’s correlation was performed and *p*-value is indicated in the plot. **C-E**, Boxplots showing the percentage of GC content (**C**), the distribution of transcript length (**D**), and the distribution of the change in translation efficiency upon sodium arsenite stress (**E**), for stress granule-enriched, -depleted or neither transcripts. **F**, Volcano plot showing differentially translated genes before and after sodium arsenite stress. **G**, Boxplots showing the distribution of translation efficiency in unstressed Hela cells, Hela cells that are treated with sodium arsenite, treated cells that are recovered for 1 hour, and treated cells that are recovered for 2 hours. Statistical analysis of **C,D,E,G** was done using Mann-Whitney U Test, with p-values labelled in the panels.

**Supplementary Figure 4. Reactivity along stress-granule enriched and depleted RNAs**. **A,B,** RNA accessibility at nucleotide resolution for NORAD (**A**) and GAPDH RNA (**B**) in unstressed (blue) and sodium arsenite stressed cells (orange).

**Supplementary Figure 5. Probing RNA structure in the stress granule deficient cells. A**, Western blot analysis showing the stabilization of USP10-peptide-DD-eGFP by treating cells with Shield-1 in the absence or presence of sodium arsenite stress. **B**, Quantification of the number of stress granules formed in unstressed cells, stressed cells and stress cells with shield-1. **C**, AUC-ROC plots of the icSHAPE reactivity based on the known accessibility of 28S rRNA.

**Supplementary Figure 6. Probing RNA-RNA interaction using SPLASH. A**, Stacked barplots showing the number and percentage of inter- and intra-molecular chimeric interactions detected in each replicates. **B**, Stacked barplots showing number of genes with detected intramolecular, intermolecular or both intra- and intermolecular interactions in unstressed and sodium arsenite treated Hela cells under indicated conditions. **C**, Scatter plots showing the correlation of chimera counts of across different biological and technical replicates for unstressed and sodium arsenite stressed cells. **D**, AUC-ROC plots of chimeric interactions using the known structure of 28S rRNA as reference. **E**, RNA-RNA interaction network of stress granule enriched and depleted RNAs in Hela cells treated with sodium arsenite. **F**, Pie charts showing the distribution of intramolecular and intermolecular interactions in 5’-UTR, CDS regions, and 3’-UTRs in cells treated with sodium arsenite. Statistical analysis of **F** was done using Chi-square Test, with p-value labelled in the panel.

**Supplementary Figure 7. Localization of SRSF proteins in cells treated with arsenite stress. A**, Immuno-fluorescence staining of G3BP1-GFP (green), SRSF1 (red), and nuclei (labeled as DAPI in blue) in Hela cells, in the presence and absence of arsenite stress. **B**, Immuno-fluorescence staining of G3BP1-GFP (green), SRSF9 (red), and nuclei (labeled as DAPI in blue) in Hela cells, in the presence and absence of sodium arsenite stress.

**Supplementary Figure 8. eCLIP analysis of SRSF1 in response to sodium arsenite stress. A**, Scatter plots showing the correlation between two biological replicates of eCLIP peaks in unstressed and sodium arsenite stressed cells. **B**, Motif enrichment of the in-house SRSF1 eCLIP data and the ENCODE SRSF1 eCLIP data. **C**, Barplots showing the number of eCLIP peaks that fall in different regions of the genome. **D**, Top, Metagene analysis of the reactivity in windows that contain SRSF1 motifs and eCLIP binding sites (blue) and in windows that only contain SRSF1 motifs but without eCLIP binding evidence (grey) in unstressed cells. Bottom, per base p-value significance of the difference between the blue and grey lines. **E**, Top, Metagene analysis of the reactivity in windows that contain SRSF1 motifs and eCLIP binding sites (red) and in windows that only contain SRSF1 motifs but without eCLIP binding evidence (grey) in sodium arsenite-stressed cells. Bottom, per base p-value significance of the difference between the blue and grey lines. **F**, Boxplots showing transcript reactivity in transcripts with low and high SRSF1 binding sites, grouped by transcript length. **G**, Boxplots showing stress granule enrichment of the transcripts with high and low SRSF1 binding sites, grouped by different transcript length. Statistical analysis of **F,G** was done using Mann-Whitney U Test, with p-values labelled in the panels. **H**, Barplots showing the amount of cell growth 48 hours after sodium arsenite treatment, in the presence and absence of SRSF1. Cell growth was measured 48 hours post stress treatment when knocking down SRSF1 in the presence and absence of SRSF1. Statistical analysis of **H** was performed using the Student’s T-Test. Data are shown as mean ± SEM, n = 5. \**p* < 0.05, \*\**p* < 0.01, \*\*\**p* < 0.001, \*\*\*\**p* < 0.0001

**Supplementary Figure 9. RNA helicases unwind RNA structures in stress granules. A**, Scatter plots showing the correlation of RT stop sites between biological replicates of indicated treatments. **B**, Barplots showing the normalized expression of DDX3X before and after siRNA knockdown. **C**, Scatter plots showing the correlation of RT stop sites between biological replicates of control and DDX3X knockdown experiments.

## Supplementary tables

**Supplementary Table 1 icSHAPE Statistics**

**Supplementary Table 2 SPLASH Statistics**

**Supplementary Table 3 SPLASH Intramolecular Interactions**

**Supplementary Table 3 SPLASH Intramolecular Interactions**

**Supplementary Table 5 oligos sequence**

