## Supplemental Figures for "RNA Structure Directs RNA Partitioning and is Actively Disrupted inside Stress Granules to Enable Cellular Recovery"

### Supplementary Figure 1

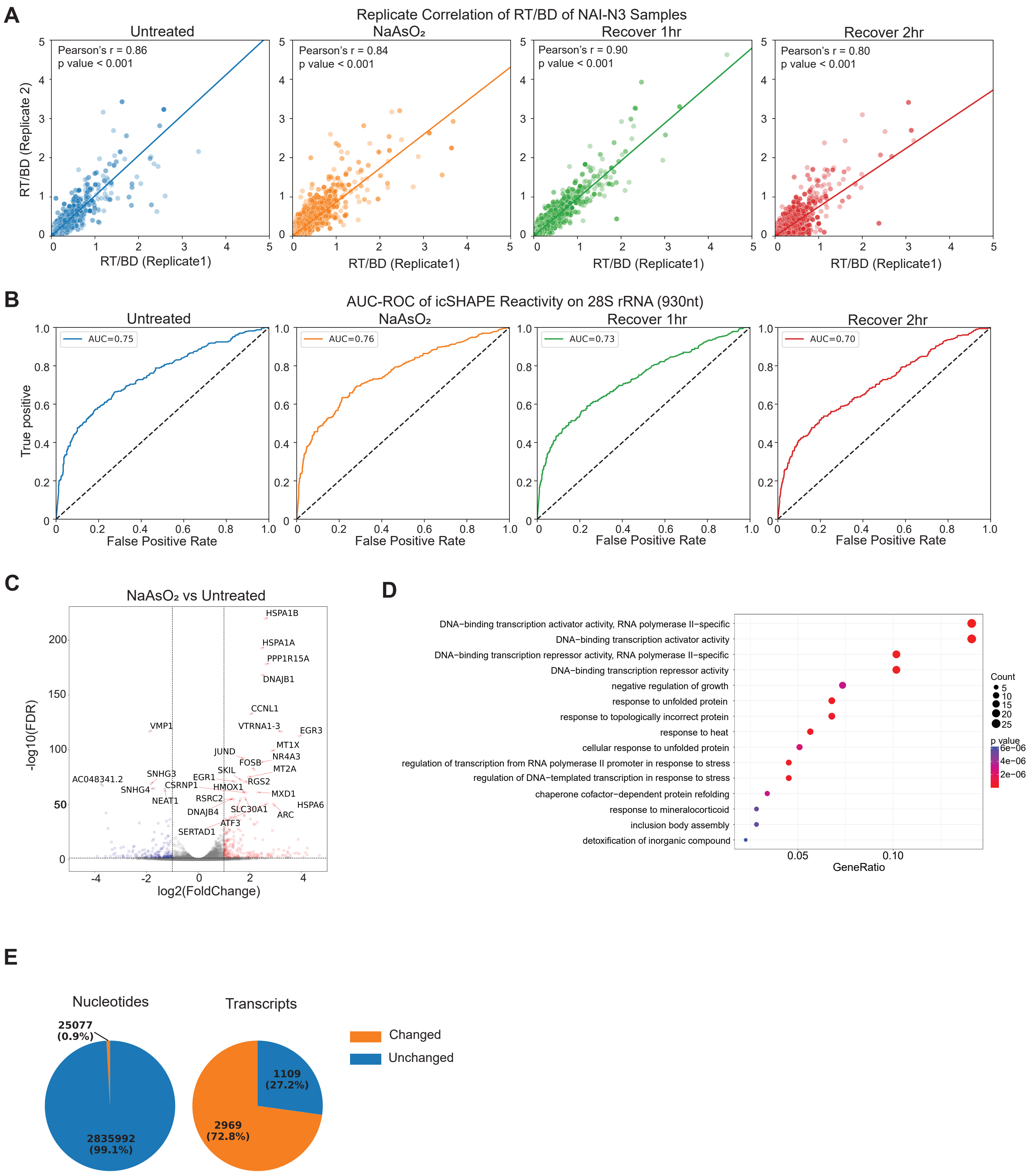

### Supplementary Figure 2

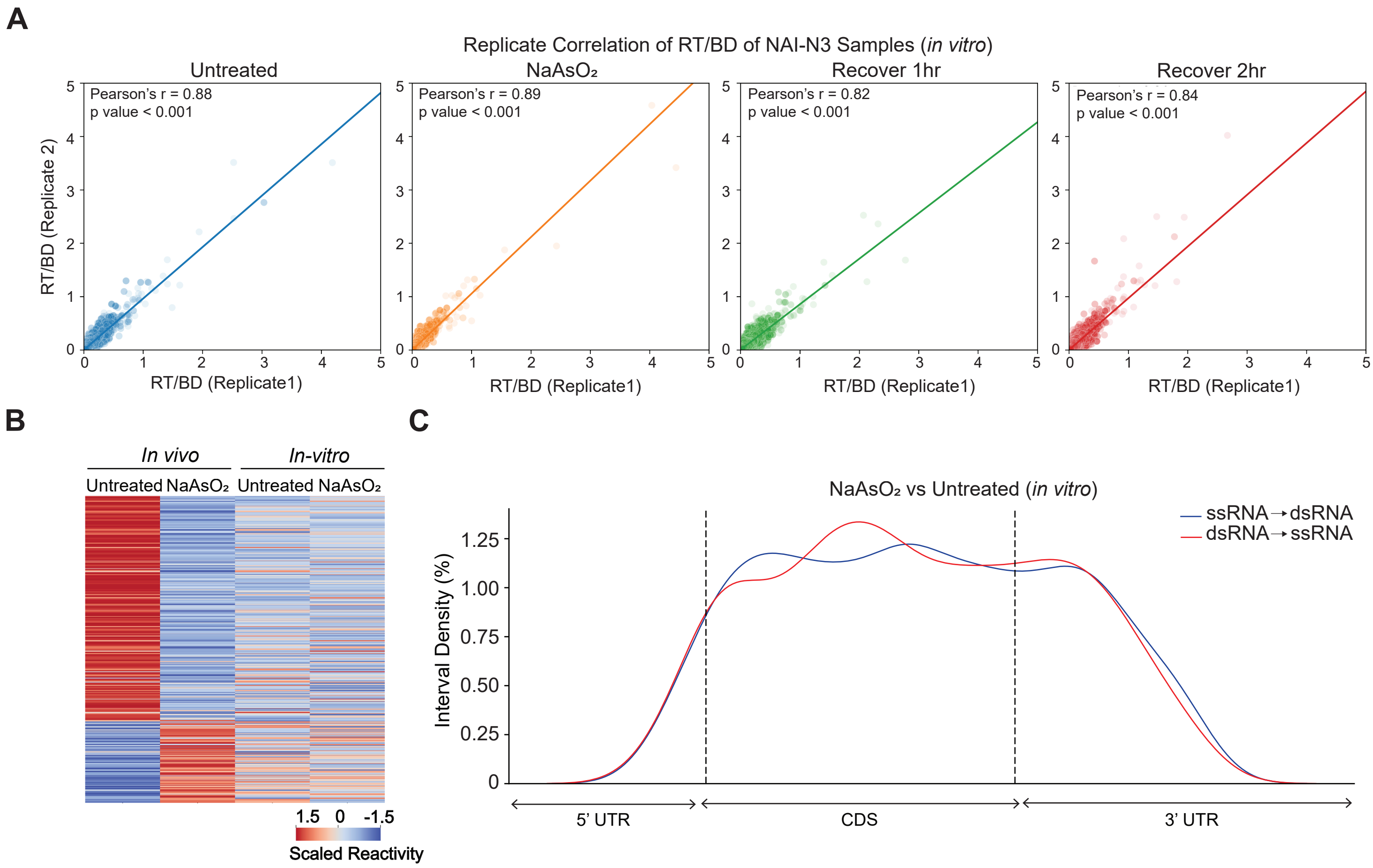

### Supplementary Figure 3

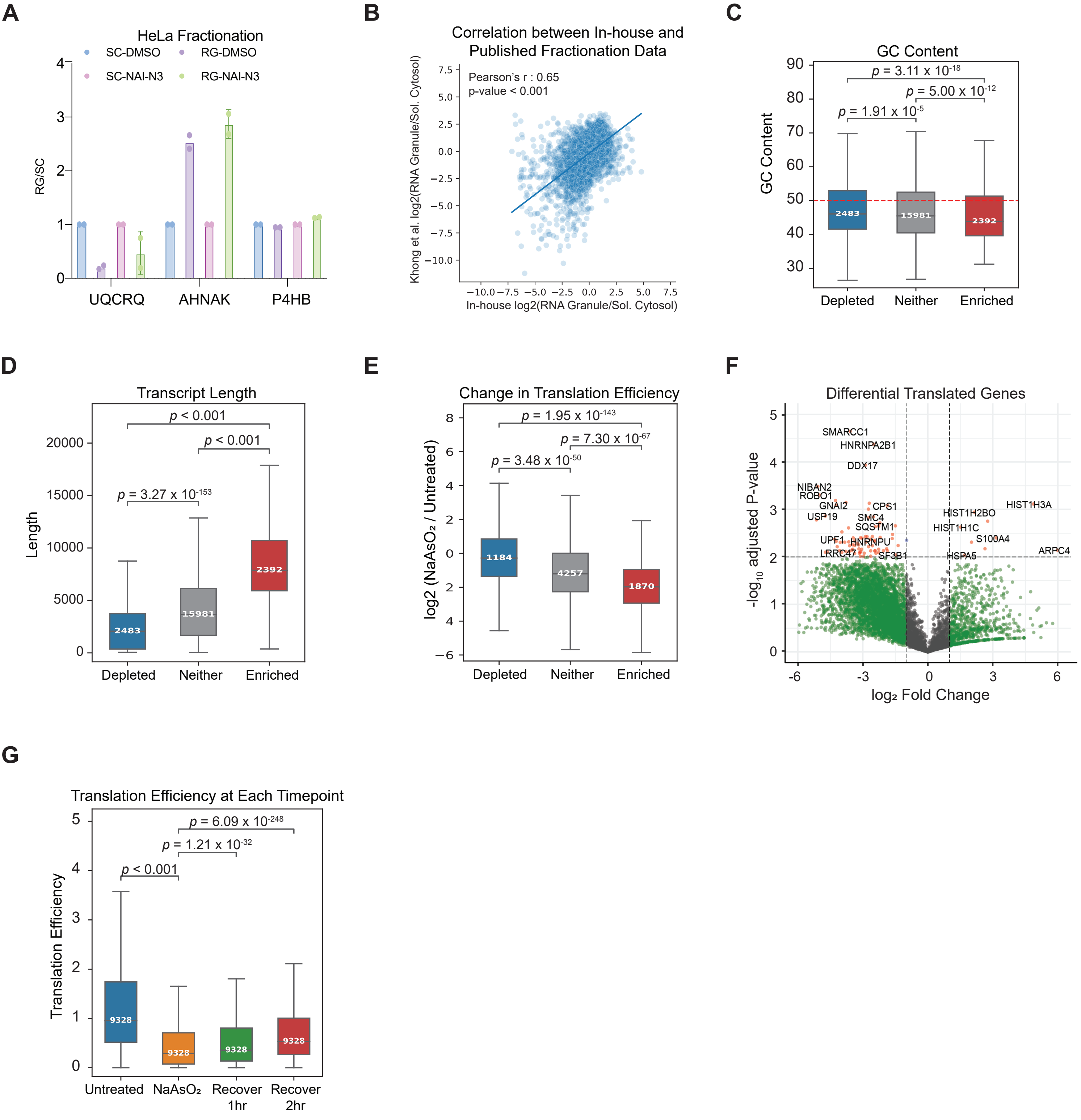

### Supplementary Figure 4

A

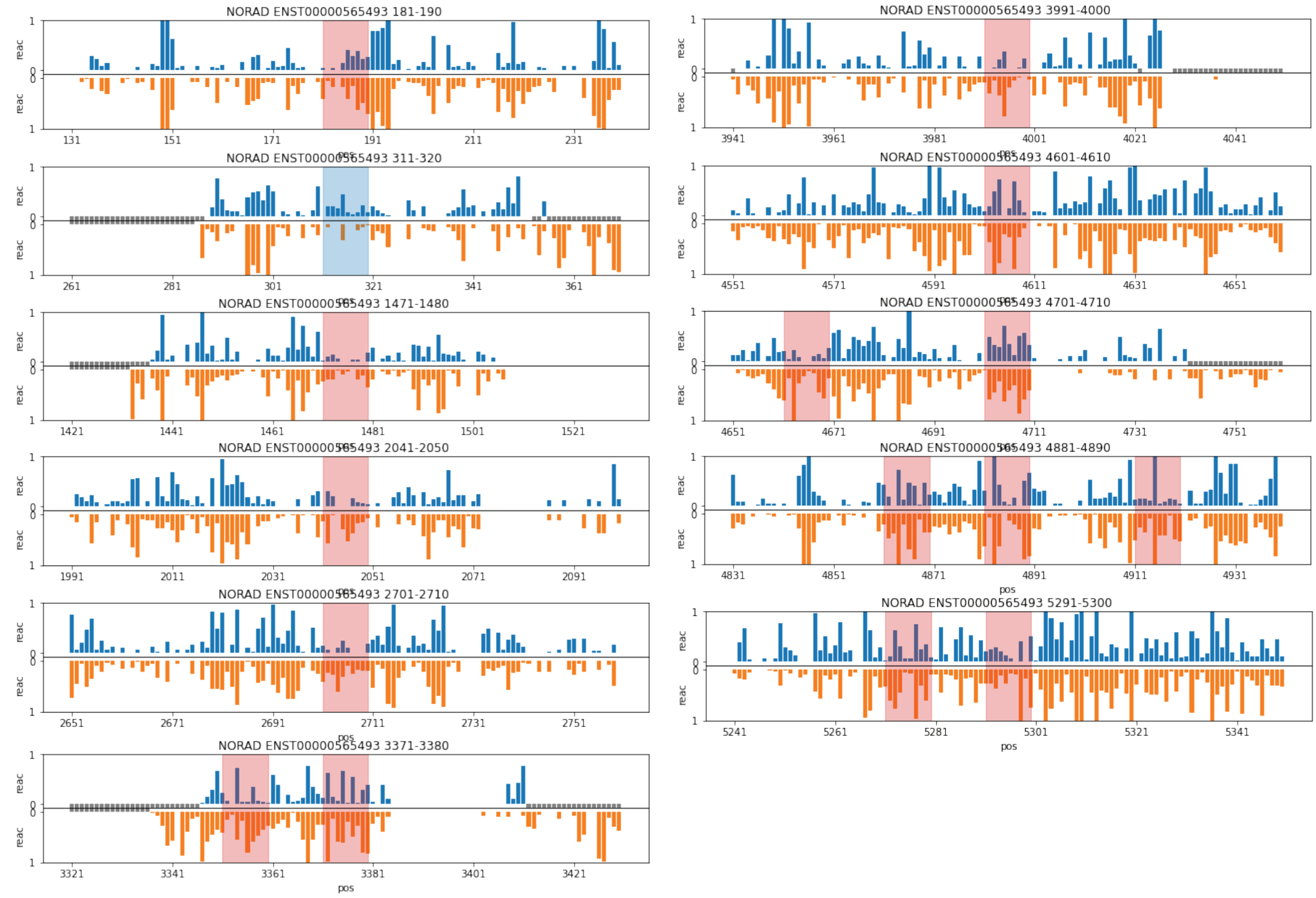


B

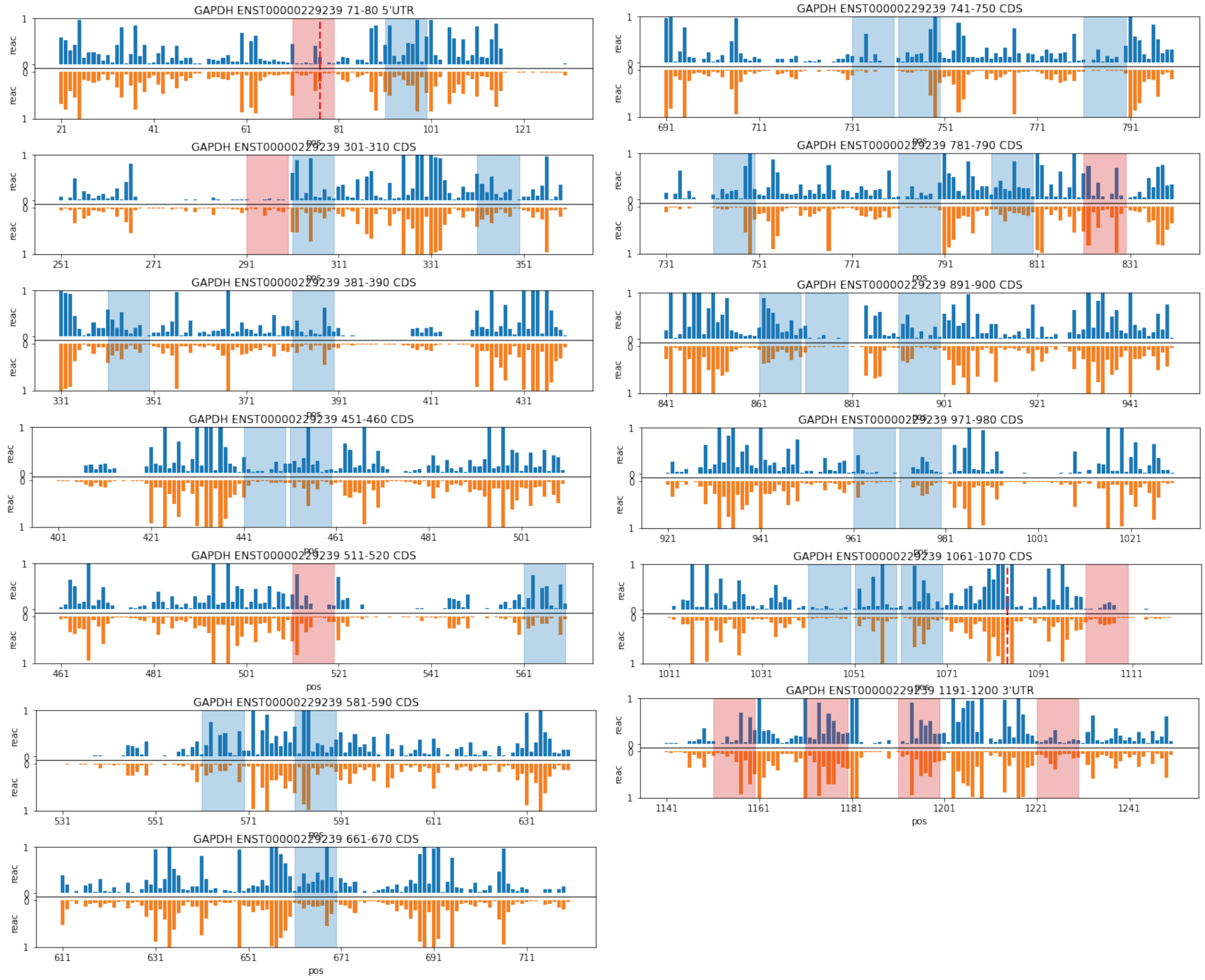

### Supplementary Figure 5

A

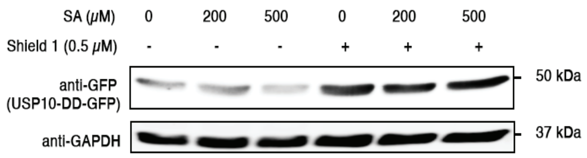


B

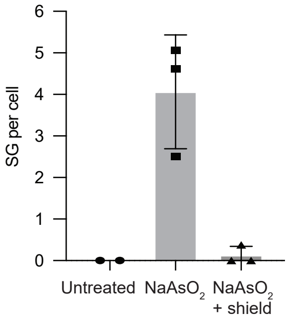


C

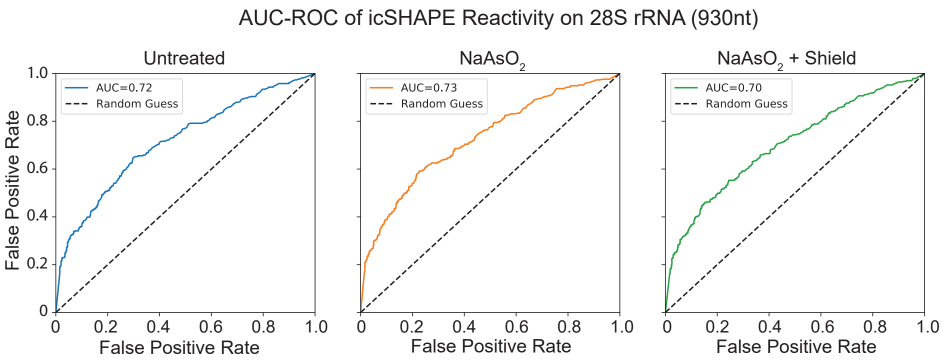

### Supplementary Figure 6

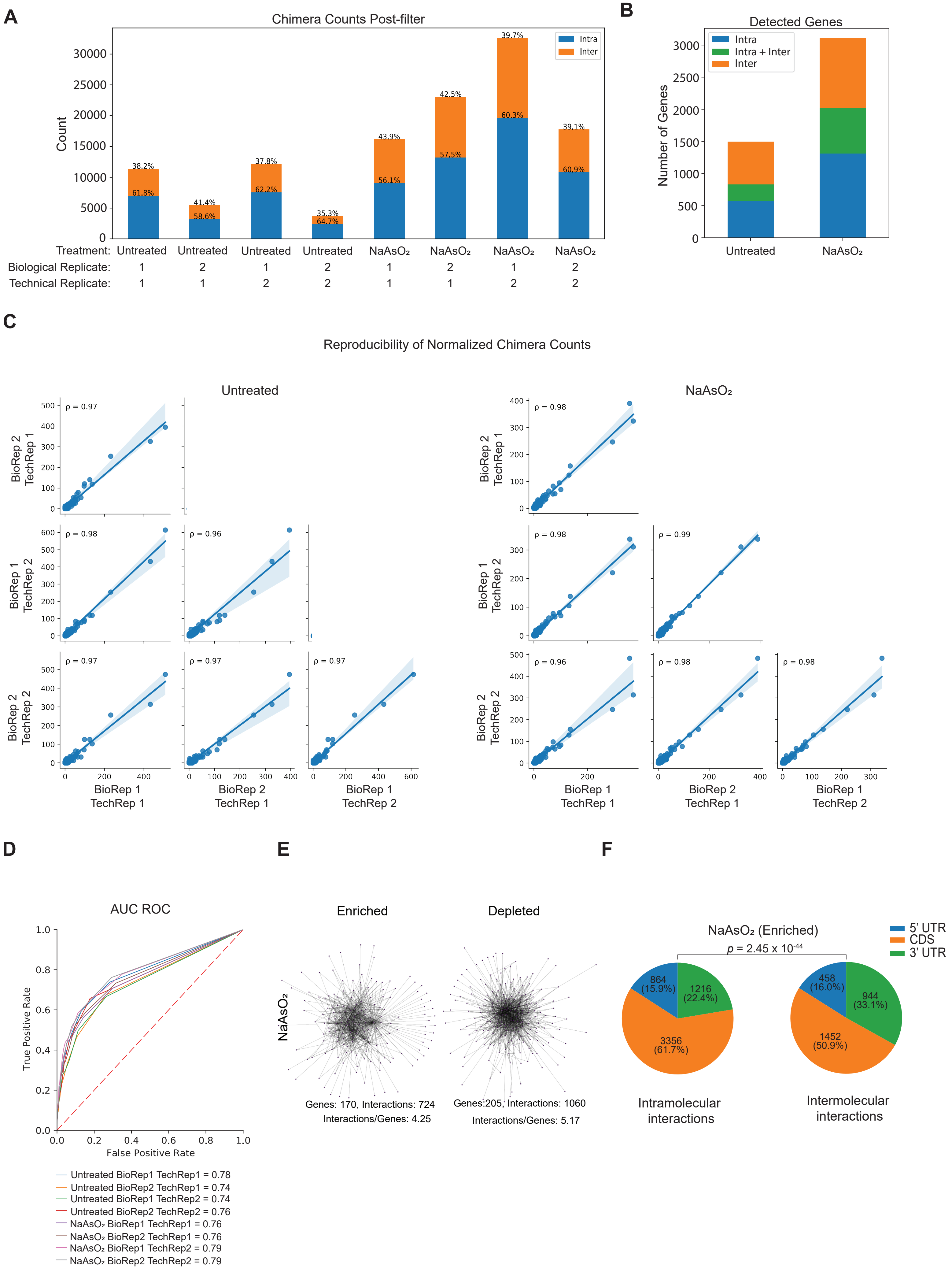

#### Supplementary Figure 7

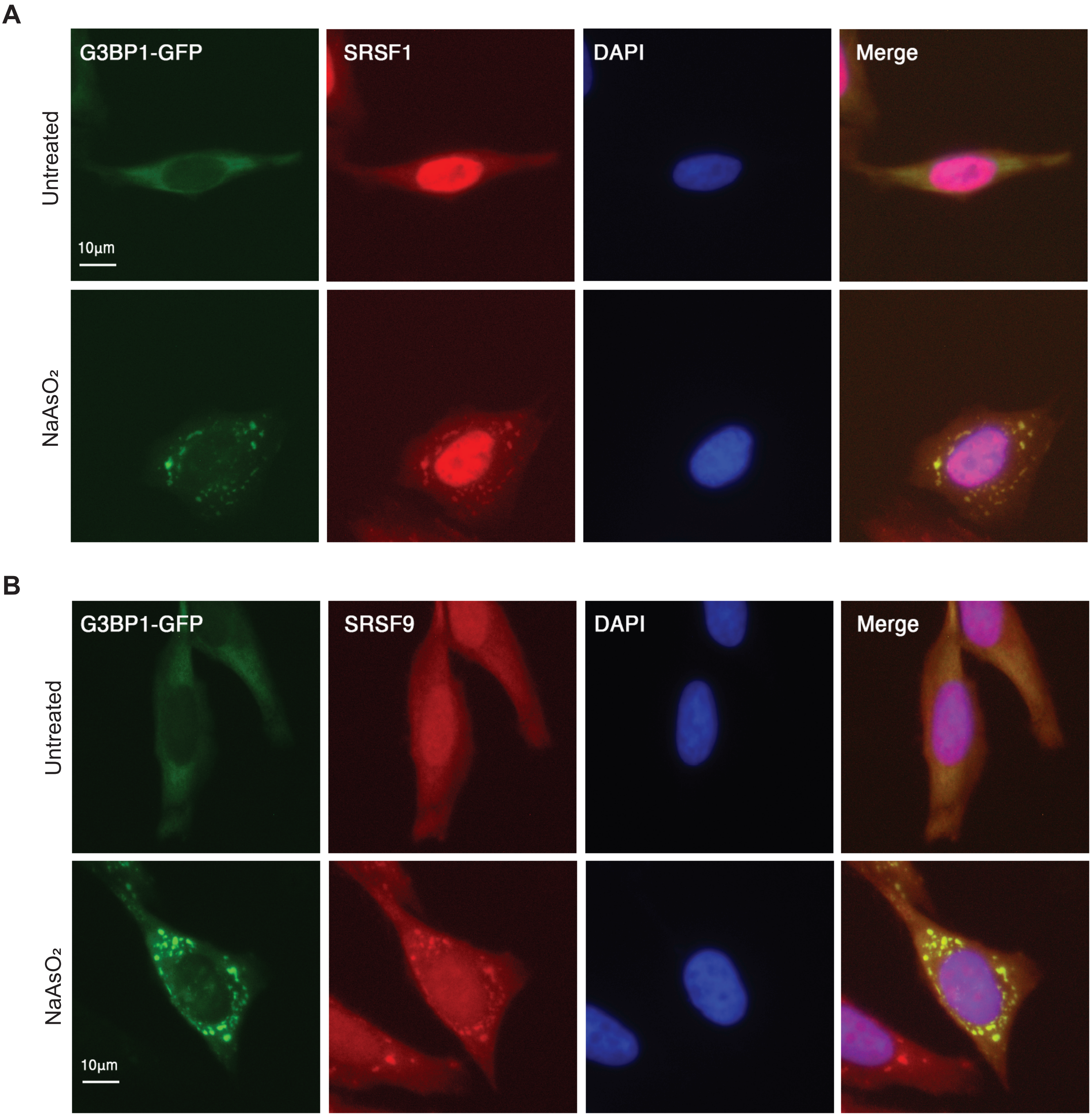

#### Supplementary Figure 8

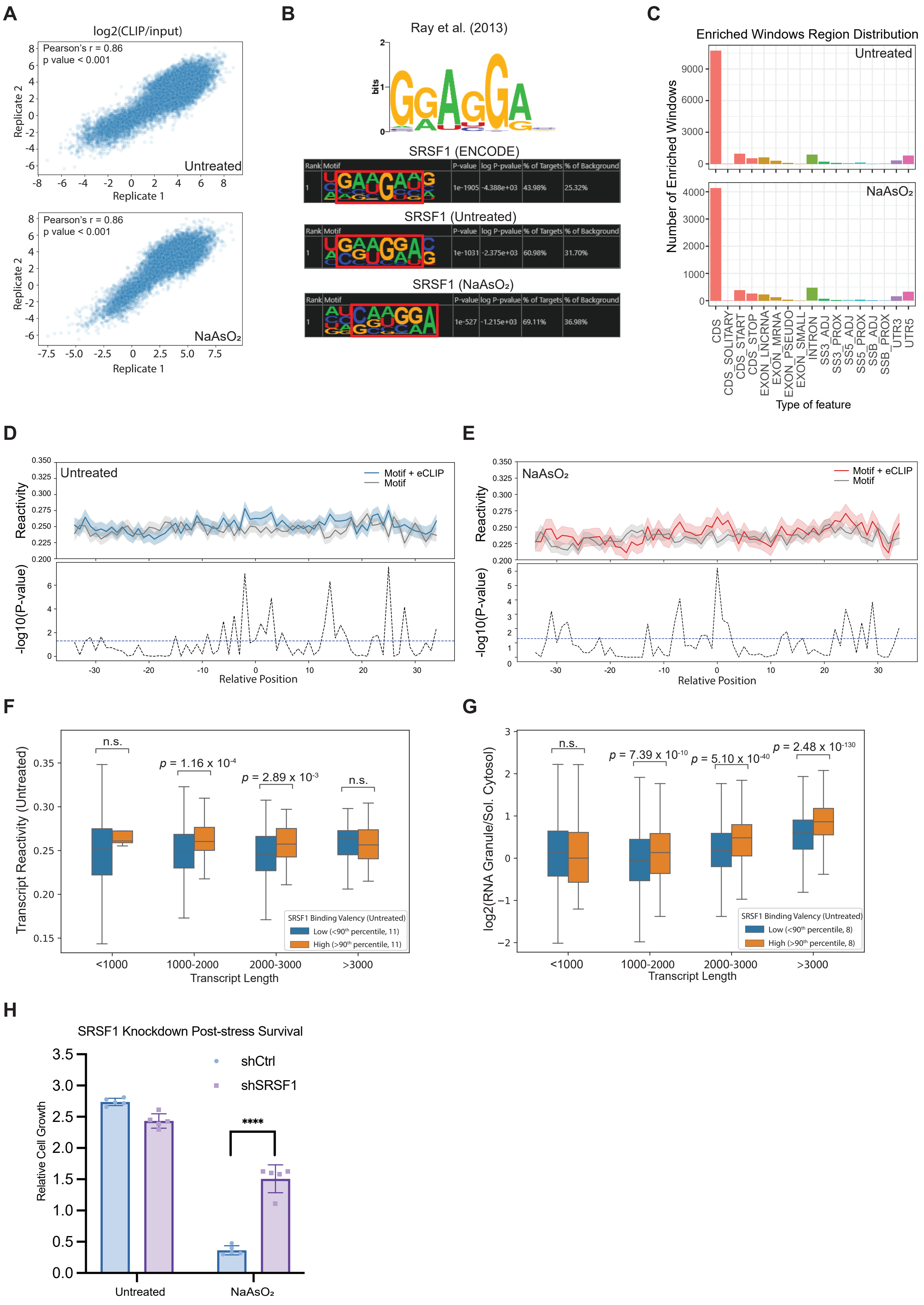

### Supplementary Figure 9

**A**

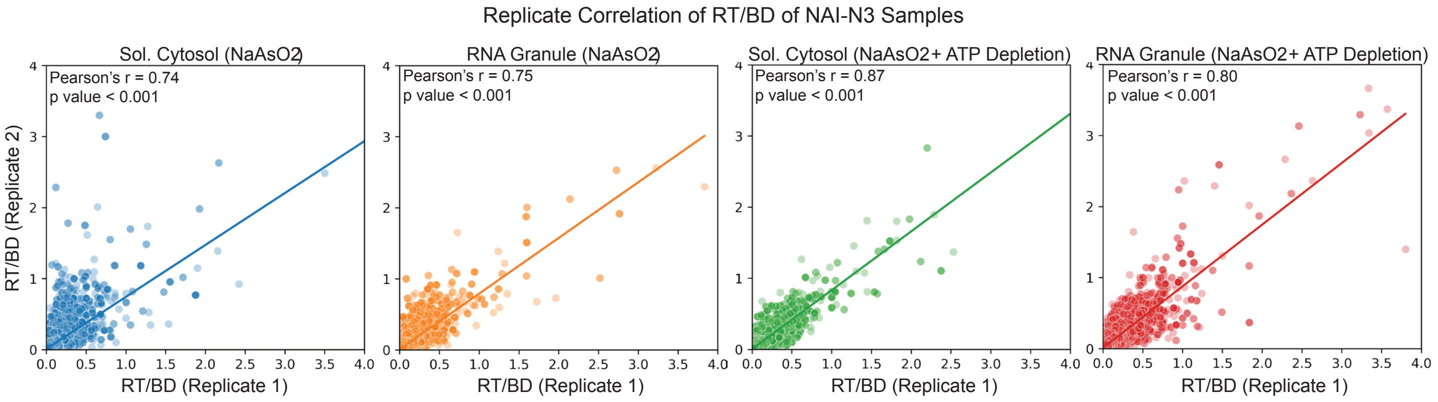


**B**

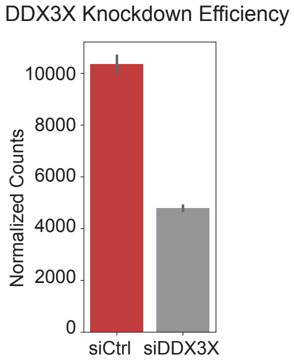


**C**

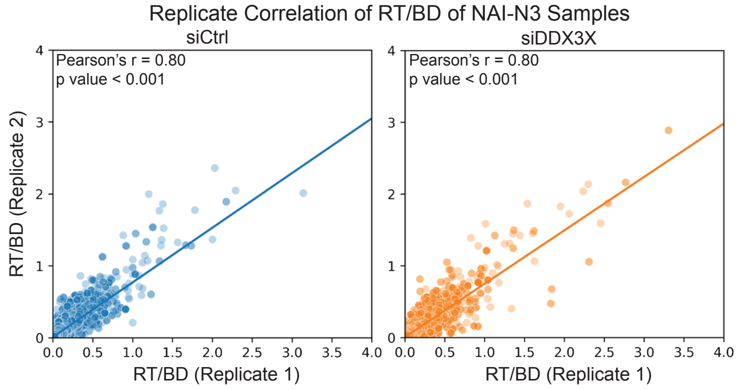
